# A potent and selective reaction hijacking inhibitor of *Plasmodium falciparum* tyrosine tRNA synthetase exhibits single dose oral efficacy *in vivo*

**DOI:** 10.1101/2024.07.22.604682

**Authors:** Stanley C. Xie, Chia-Wei Tai, Craig J. Morton, Liting Ma, Shih-Chung Huang, Sergio Wittlin, Yawei Du, Yongbo Hu, Con Dogovski, Mina Salimimarand, Robert Griffin, Dylan England, Elisa de la Cruz, Ioanna Deni, Tomas Yeo, Anna Y. Burkhard, Josefine Striepen, Kyra A. Schindler, Benigno Crespo, Francisco J. Gamo, Yogesh Khandokar, Craig A. Hutton, Tayla Rabie, Lyn-Marié Birkholtz, Mufuliat T. Famodimu, Michael J. Delves, Judith Bolsher, Karin M. J. Koolen, Rianne van der Laak, Anna C. C. Aguiar, Dhelio B. Pereira, Rafael V. C. Guido, Darren J. Creek, David A. Fidock, Lawrence R. Dick, Stephen L. Brand, Alexandra E. Gould, Steven Langston, Michael D.W. Griffin, Leann Tilley

## Abstract

The *Plasmodium falciparum* cytoplasmic tyrosine tRNA synthetase (*Pf*TyrRS) is an attractive drug target that is susceptible to reaction-hijacking by AMP-mimicking nucleoside sulfamates. We previously identified an exemplar pyrazolopyrimidine ribose sulfamate, ML901, as a potent pro-inhibitor of *Pf*TyrRS. Here we examined the stage specificity of action of ML901, showing very good activity against the schizont stage, but lower trophozoite stage activity. We explored a series of ML901 analogues and identified ML471, which exhibits improved potency against trophozoites and enhanced selectivity against a human cell line. Additionally, it has no inhibitory activity against human ubiquitin-activating enzyme (UAE) *in vitro*. ML471 exhibits low nanomolar activity against asexual blood stage *P. falciparum* and potent activity against liver stage parasites, gametocytes and transmissible gametes. It is fast-acting and exhibits a long *in vivo* half-life. ML471 is well-tolerated and shows single dose oral efficacy in the SCID mouse model of *P. falciparum* malaria. We confirm that ML471 is a pro-inhibitor that is converted into a tight binding Tyr-ML471 conjugate by the *Pf*TyrRS enzyme. A crystal structure of the *Pf*TyrRS/ Tyr-ML471 complex offers insights into improved potency, while molecular docking into UAE provides a rationale for improved selectivity.

## Introduction

Malaria is a debilitating disease caused by protist parasites of the genus *Plasmodium* that places an enormous health burden on the world’s poorest communities. In 2022, more than 200 million people were infected with *P. falciparum*, resulting in more than 600,000 deaths [1]. The burden was exacerbated by disruptions to services during the COVID pandemic [2]. Unfortunately, the past 15 years have seen the emergence of *P. falciparum* parasites that exhibit partial resistance to artemisinin and partner drugs, such as piperaquine and mefloquine, resulting in ∼50% treatment failure with standard artemisinin combination therapies in some regions of Southeast Asia [3, 4]. The recent emergence in Africa of parasites harbouring artemisinin resistance-conferring K13 mutations [5–7] is of great concern, and the Medicines for Malaria Venture (MMV) not-for-profit partnership has declared that new antimalarial therapies and prophylaxis regimens need to be developed as a failsafe [8].

Certain AMP-mimicking nucleoside sulfamates act as pro-inhibitors of E1 enzymes, *i.e.* ubiquitin/ ubiquitin-like protein (UBL)-activating enzymes [9–13]. The UBL-bound form of these Adenylate-Forming Enzymes (AFEs) is susceptible to attack, leading to the formation of an inhibitory sulfamate-UBL adduct within the active site. This unusual reaction hijacking mechanism has been exploited to generate new clinical candidates, such as Pevonedistat [14, 15].

We previously screened a Takeda Pharmaceuticals nucleoside sulfamates library (Cambridge, MA, USA) and identified ML901 as an exemplar pyrazolopyrimidine sulfamate, with potent activity against *P. falciparum* [16]. Our group showed that, surprisingly, this AMP-mimicking nucleoside sulfamate uses a related reaction hijacking mechanism to target another AFE subclass, the amino acyl tRNA synthetases. ML901 binds the *P. falciparum* cytoplasmic tyrosine tRNA synthetase (*Pf*TyrRS), and then reacts with the bound activated amino acid, resulting in the synthesis of an inhibitory sulfamate-amino acid adduct within the active site of the enzyme. By contrast, the equivalent human enzyme is not susceptible to reaction hijacking. That finding was the first demonstration of reaction hijacking of an enzyme class other than the E1 enzymes. More recently, we have identified an aminothienopyrimidine sulfonamide, OSM-S-106, with selective reaction hijacking activity against the *P. falciparum* asparagine tRNA synthetase [17].

ML901 exhibits low, but measurable toxicity against a mammalian cell line [16], which is potentially due to cross-inhibition of UBLs [12]. Thus, we explored a range of ML901 derivatives from the Takeda nucleoside sulfamate library, with different substitutions at the 7-position of the pyrazolopyrimidine ring system, to identify compounds with improved selectivity. We identified ML471 as a compound with enhanced potency against *P. falciparum* and decreased activity against human ubiquitin-activating enzyme (UAE). ML471 exhibits enhanced cellular and biochemical selectivity. It also exhibits activity against plasmodium liver stages and sexual transmissible stages. Importantly, ML471 exhibits rapid killing kinetics and demonstrates single dose oral efficacy against *P. falciparum* in an *in vivo* model.

## Results and Discussion

### Potency and selectivity of ML901 derivatives

We examined the activity of a series of pyrazolopyrimidine sulfamates with different substitutions at the 7-position (Fig. 1A-I), from the Takeda Pharmaceuticals Library, against the growth of asexual blood stage *P. falciparum* (3D7 strain) in a 72-h exposure assay. Consistent with our previous report [16], ML901 exhibits potent activity with a 50% Inhibitory Concentration (IC_50_72h_) of 2.8 nM (Table 1). Indeed, most of the compounds show excellent potency (Table 1), demonstrating that different substitutions at the 7-position are well-tolerated, except for the bulky phenoxy substituent (ML470) (IC_50_72h_ = 44.8 nM; Table 1). The non-specific inhibitor, adenosine 5’-monosulfamate (AMS, Fig. 1J [16]), which has a different heterocyclic base and lacks the 7-position substituent, also exhibits potent activity (IC_50_ = 1.8 nM; Table 1). ML471, which bears an isopropyl group at the 7-position, exhibits very potent activity (IC_50_72h_ = 1.5 nM, Table 1).

**Figure 1.**
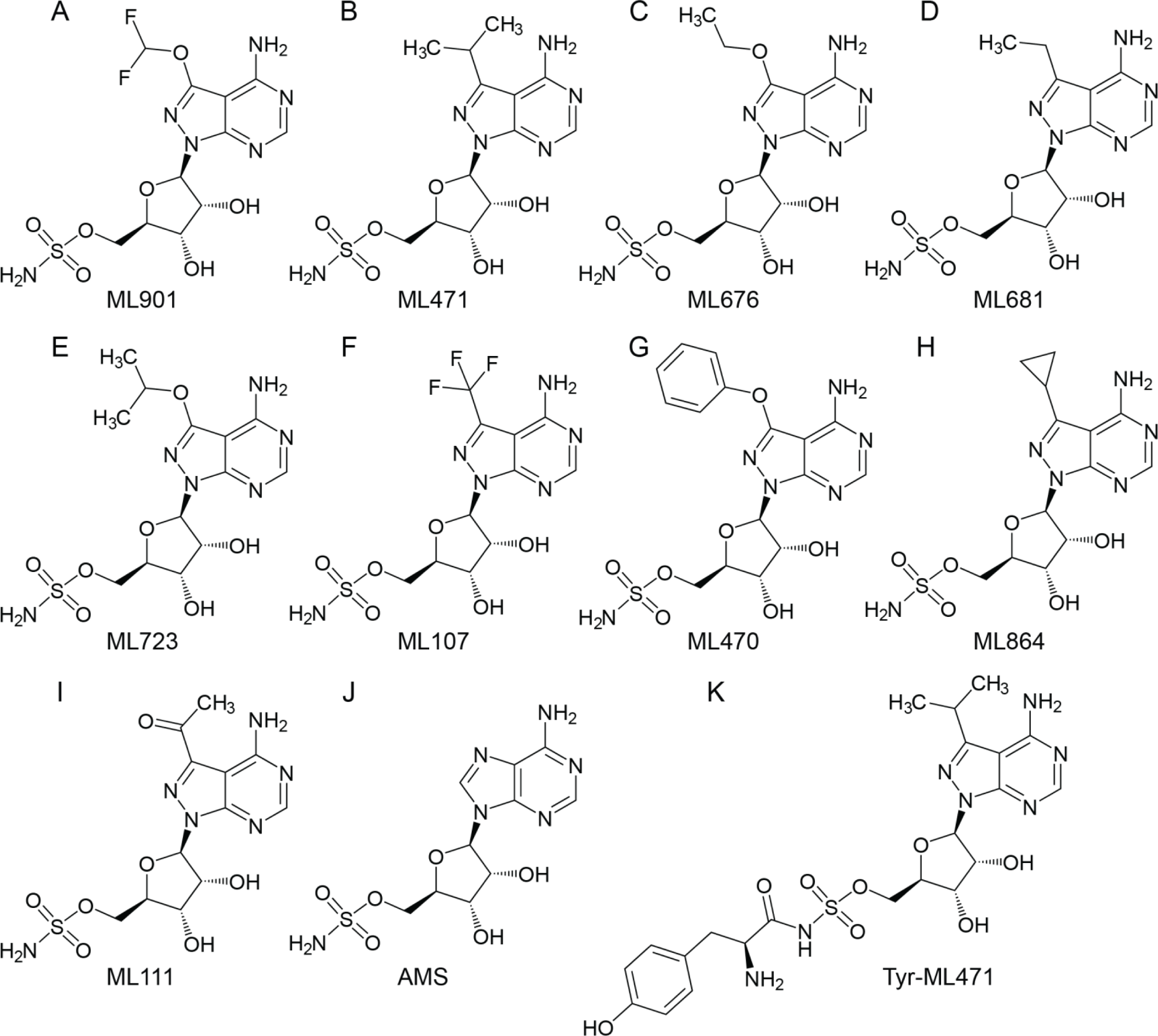
Structures of ML901 and derivatives and adenosine 5’-sulfamate (AMS). (A) ML901, (B) ML471, (C) ML676, (D) ML681, (E) ML723, (F) ML107, (G) ML470, (H) ML864, (I) ML111, (J) AMS, (K) Tyr-ML471.

**Table 1.** Activities of pyrazolopyrimidine sulfamates as inhibitors of parasite growth. AMS = Adenosine 5’-sulfamate. n = Number of biological repeats. Data values represent mean ± SEM. Medicines for Malaria Venture (MMV) designations are in brackets.

| Compound | *P. falciparum* (3D7) |  | *H. sapiens* (HepG2) |
| --- | --- | --- | --- |
|  | 72-h IC<sub>50</sub> (nM) | 6-h Trophozoite Stage IC<sub>50</sub> (nM) | 72-h IC<sub>50</sub> (nM) |
| **ML901**<br>(MMV1581329) | 2.8 ± 0.2 (n = 3) | 135 ± 14 (n = 3) | 4,650 ± 1,390 (n = 3) |
| **ML471**<br>(MMV1793207) | 1.5 ± 0.2 (n = 3) | 29.1 ± 3.0 (n = 3) | > 50,000 (n = 3) |
| **ML676**<br>(MMV1793313) | 2.8 ± 0.2 (n = 3) | N/A | 22,400 ± 7,300(n = 3) |
| **ML681**<br>(MMV1793314) | 3.4 ± 0.1(n = 3) | N/A | 7,530 ± 1,250 (n = 3) |
| **ML723**<br>(MMV1793208) | 2.5 ± 0.6 (n = 3) | 220 ± 31 (n = 3) | > 50,000 (n = 3) |
| **ML107**<br>(MMV1793318) | 4.2 ± 0.5 (n = 3) | 150 ± 33 (n = 3) | 2,520 ± 820 (n = 3) |
| **ML470**<br>(MMV1793342) | 44.8 ± 8.2 (n = 4) | N/A | > 50,000 (n = 3) |
| **ML864**<br>(MMV1793301) | 2.5 ± 0.6 (n = 3) | N/A | 21,300 ± 1,700 (n = 3) |
| **ML111**<br>(MMV1793328) | 4.3 ± 0.7 (n = 3) | N/A | > 50,000 (n = 3) |
| **AMS** | 1.8 ± 0.6 (n = 3)* | N/A | N/A |

\* Data from [16]

Short duration pulse exposure to antimalarial drugs can be used to dissect differences in potency of compounds against different stages of development that may not be evident in standard 72-h exposure assays [18]. Here we subjected tightly synchronised blood stage *P. falciparum* cultures to different duration pulses of ML901 and measured parasitemia in the next cycle [19]. Schizont stage parasites are efficiently killed by ML901, even when exposed to pulses as short as 3 h or 6 h (Supplementary Fig. 1B, Table S1). This may reflect the need for synthesis of daughter merozoite proteins during schizogony. By contrast, trophozoite stage parasites are 10 to 20-fold less sensitive (Supplementary Fig. 1A, Table S1).

To determine whether ML471 and other selected pyrazolopyrimidine sulfamates exhibit enhanced potency, we exposed synchronised trophozoite stage cultures to 6-h pulses and measured growth inhibition by quantifying the SYBR Green I fluorescence signal in the next cycle. ML471 exhibits enhanced potency (IC_50_6h_ = 29.1 nM, Fig. 2A, Table 1) compared with ML901 and the other analogues tested (Fig. 2A, Table 1, IC_50_6h_ values ranging from 135 nM to 220 nM).

**Figure 2.**
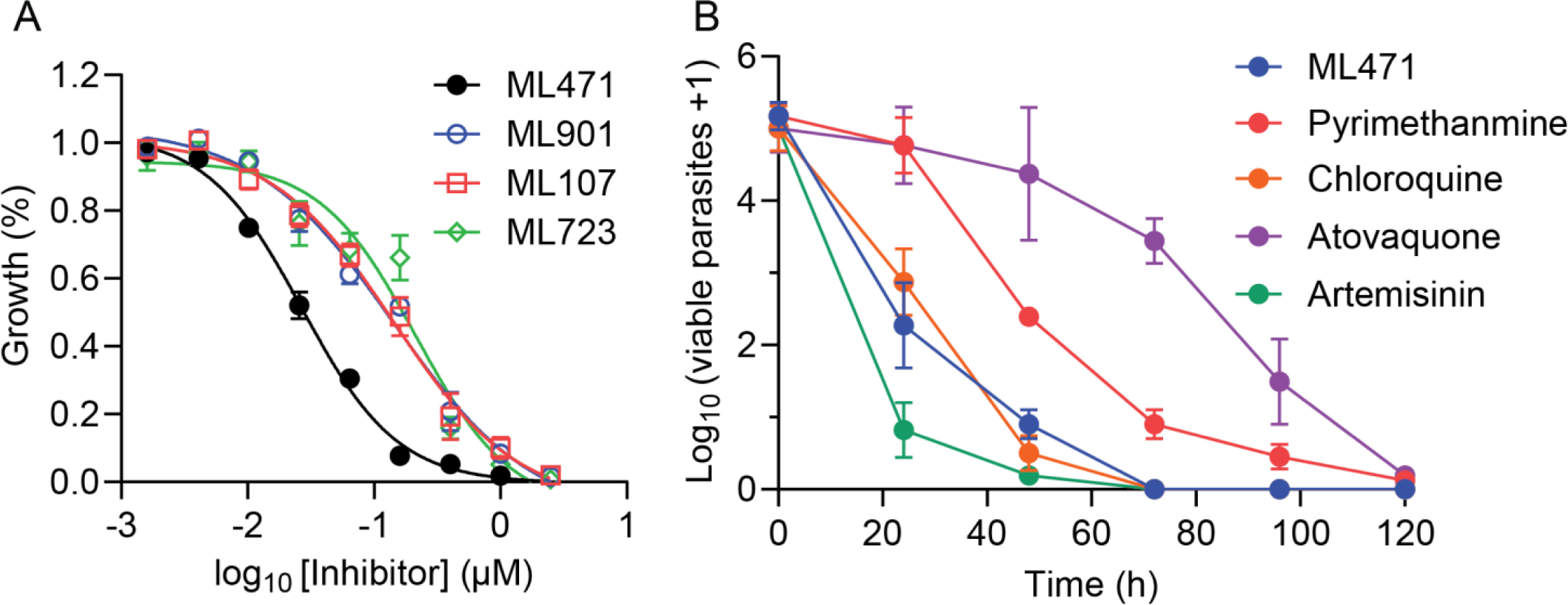
ML471 exhibits improved short-exposure activity against *P. falciparum* cultures, associated with rapid parasite killing. (A) Synchronized Cam3.II^rev^ parasite cultures were subjected to 6-h pulses of ML901, ML471, ML107 and ML723, at the trophozoite (25-30 h.p.i.) stage. Growth inhibition was determined in the cycle following treatment. Data represent the mean of three independent experiments and error bars correspond to SEM. (B) 3D7 parasite cultures were treated for 0 to 120 h with ML471 or compounds with fast (artemisinin, chloroquine), moderate (pyrimethamine) or slow (atovaquone) killing profiles, at 10 times their respective IC_50_48h_ values. Following removal of inhibitor, serial dilutions of cultures were established, and assessed after 18 days of culturing.

We examined the toxicity of the compounds against the human HepG2 cell line, employing a 72-h exposure period. ML901 inhibits the growth of HepG2 cells with an IC_50_72h_ of 4.65 μM (Table 1). Some of the compounds from the pyrazolopyrimidine sulfamate series, including ML471 exhibited markedly improved selectivity, with IC_50_ values above the range of the assay (>50 μM, Table 1). Other compounds such as ML107, which has a trifluoromethyl substituent, and ML681 and ML864 which have slightly smaller substituents as compared to an isopropyl group, show a higher level of toxicity against the mammalian cell line. These data suggest that the size of the 7-position substituent impacts selectivity. Our previous report [16] showed that AMS is also toxic to mammalian cells lines, such as HCT116 (IC_50_72h_ = 26 nM), in agreement with previous reports [20, 21], and, as expected, given its broad inhibitory activity.

### ML471 exhibits low activity against critical E1 enzymes

ML901 was originally investigated as an inhibitor of Atg7, an E1 that activates the Autophagy-related protein 8 (Atg8), a ubiquitin-like protein involved in the formation of autophagosomal membranes [12, 22]. However, Atg7 is not essential for cell survival *in vitro* [23] and is unlikely to underpin the observed mammalian cell toxicity [12]. By contrast, loss-of-function of other E1 enzymes, in particular UAE, is known to be deleterious to the survival and growth of cells [10, 11, 24, 25].

To explore the molecular basis of the enhanced cellular selectivity of ML471 and other derivatives compared with ML901, we assessed inhibitory activity against a range of E1 enzymes. ML901 exhibits strong inhibition of Atg7 (IC_50_ = 33 nM) and clinically relevant activity against UAE (IC_50_ = 5.39 μM), with lower-level activity against NEDD8 Activating Enzyme (NAE IC_50_ = 28 μM) and no activity against SUMO Activating Enzyme (SAE) (Table 2). ML676, ML723, ML681, ML107 and ML111 also exhibit relevant activity against one or more of UAE, NAE or SAE, consistent with their weak to moderate cellular toxicity. Such off-target activity could limit the development of these compounds. As previously reported [16], AMS is a potent inhibitor of each of the E1s tested (Table 2), likely contributing to the high cellular toxicity. ML471 inhibits the activity of human Atg7 (IC_50_ = 22 ± 9 nM), but exhibits no or very little activity against UAE, NAE and SAE, consistent with low mammalian cell cytotoxicity.

**Table 2.** Inhibitory activity of selected pyrazolopyrimidine sulfamates in E1 enzymes assays. ATG7 = autophagy-related protein-7. NAE = NEDD8-activating enzyme. UAE = ubiquitin activating enzyme. SAE = SUMO-activating enzyme. GABARAP = GABAA receptor-associated protein. HTRF = Homogeneous Time-Resolved Fluorescence. Data represent mean ± SEM. n = Number of independent experiments.

| Compound | ATG7<br>IC<sub>50</sub> HTRF<br>RH-GABARAP (μM) | NAE<br>IC<sub>50</sub> HTRF (μM) | UAE<br>IC<sub>50</sub> HTRF (μM) | SAE<br>IC<sub>50</sub> HTRF (μM) |
| --- | --- | --- | --- | --- |
| **ML901** | 0.033 ± 0.003 (n = 57)* | 28.0 ± 0.6 (n=3) | 5.39 ± 0.160 (n = 3) | >100 (n = 3) |
| **ML471** | 0.022 ± 0.009 (n = 6) | >100 (n=3) | 85.7 ± 6.5 (n=3) | >100 (n = 3) |
| **ML681** | N.A. | 35.3 ± 0.8 (n=3) | 8.7 ± 0.6 (n=3) | 22.4 ± 3.1 (n=3) |
| **ML676** | N.A. | 66.6 ± 1.1 (n=3) | 73.4 ± 3.5 (n=3) | 31.4 ± 5.7 (n=3) |
| **ML723** | N.A. | >100 (n=3) | >100 (n=3) | >100 (n=3) |
| **ML470** | N.A. | >100 (n=3) | >100 (n=3) | >100 (n=3) |
| **ML107** | N.A. | 75.3 ± 0.8 (n=3) | 22.9 ± 0.7 (n=3) | 78.5 ± 6.2 (n=3) |
| **AMS\*** | 410 ± 20 (n = 90) | 0.006 ± 0.001 (n = 9) | 0.006 ± 0.003 (n = 3) | 0.006 ± 0.002 (n = 7) |

\*Data from [16]

### Parasitological properties of ML471

Given its enhanced potency and selectivity, ML471 was selected for further characterisation. In addition to potent and selective activity against laboratory asexual blood stage *P. falciparum,* new antimalarial compounds should exhibit activity against clinical strains of *P. falciparum* and *P. vivax* and, preferably, exhibit activity against liver and transmissible stages. ML471 exhibits potent activity against South American clinical isolates of *P. falciparum* and *P. vivax*, including chloroquine-resistant strains, with median IC_50_ values of 4.0 nM (*Pf*) and 6.7 nM (*Pv*), respectively, similar to artesunate (Table S2). ML471 exhibits improved potency compared with ML901 against gametocytes at both the early (IC_50_ = 112 nM) and mature (IC_50_ = 392 nM) stages of development, with potencies similar to those for Methylene Blue and the *Plasmodium* phosphatidylinositol 4-kinase (PI4K) inhibitor, MMV390048 (Table S3). ML471 prevents development of both *P. falciparum* NF175 and NF135 schizonts in primary human hepatocytes with high potency (IC_50_ = 2.8 nM for NF175 and IC_50_ = 5.5 nM for NF135, Table S4), while exhibiting no toxicity against the primary hepatocyte host cells (Table S4). ML471 potently inhibits the fertility of transmissible male (IC_50_ = 49 nM) and female (IC_50_ = 260 nM) gametocytes (Table S5). The positive control, Cabamiquine, exhibited potent activity against both gametocyte sexes, consistent with a previous report [26]. In each of these assays, ML471 exhibits similar or improved potency compared with ML901 (Table S3-S5).

The Parasite Reduction Rate (PRR) was assessed using a standardized method [27] and compared with compounds exhibiting very fast (artemisinin), fast (chloroquine), moderate (pyrimethamine) or slow (atovaquone) killing profiles, at 10 times their respective IC_50_48h_ values. The Log PRR for ML471 of 4.1 is considered fast, and is similar to chloroquine (Fig. 2B, Table S6).

### Pharmacological properties of ML471

To meet MMV candidate selection criteria new antimalarial compounds for treatment indications would minimally need to have an oral dose of <500 mg to achieve a 12-log kill in a 55 kg adult. ML471 exhibits a favourably low molecular weight (MW = 388) and good solubility (Table S7). It has a predicted AlogP of −1.18 and a Topological Polar Surface Area (TSPA) of 189 Å^2^, which suggest this compound may have difficulty being absorbed but should be metabolically stable. As expected, rat oral bioavailability needs optimisation (%F (blood) = 8.72 to 9.56, n = 3) and renal clearance of the parent compound was evident (9-38% of the dose recovered in 0-24 h urine as parent). Despite the sub-optimal oral absorption, the rat pharmacokinetic profile of ML471 (25 mg/kg p.o.; Fig. 3B; Table S7) exhibits excellent duration of absorbed drug exposure. The area under the curve is 30 μM.h, reflecting the low blood clearance (∼4% of liver blood flow after an IV dose of 1 mg/kg) and the long terminal half-life in blood (T_1/2∞_ = 30.5 h) (Table S7). Acting in its favour, ML471 shows high retention in red blood cells (RBCs) in the i.v. PK study, with blood to plasma ratios around 1 at the initial sampling times, but increasing greatly over time due to slower clearance from the RBC compartment (Table S7; and compare Fig. 3A and B). ML471 contains a sulfamate group that is predicted to bind tightly to carbonic anhydrase [28, 29], which likely explains accumulation of ML471 into RBCs (where carbonic anhydrase is abundant and where the asexual stage parasites are located).

**Figure 3.**
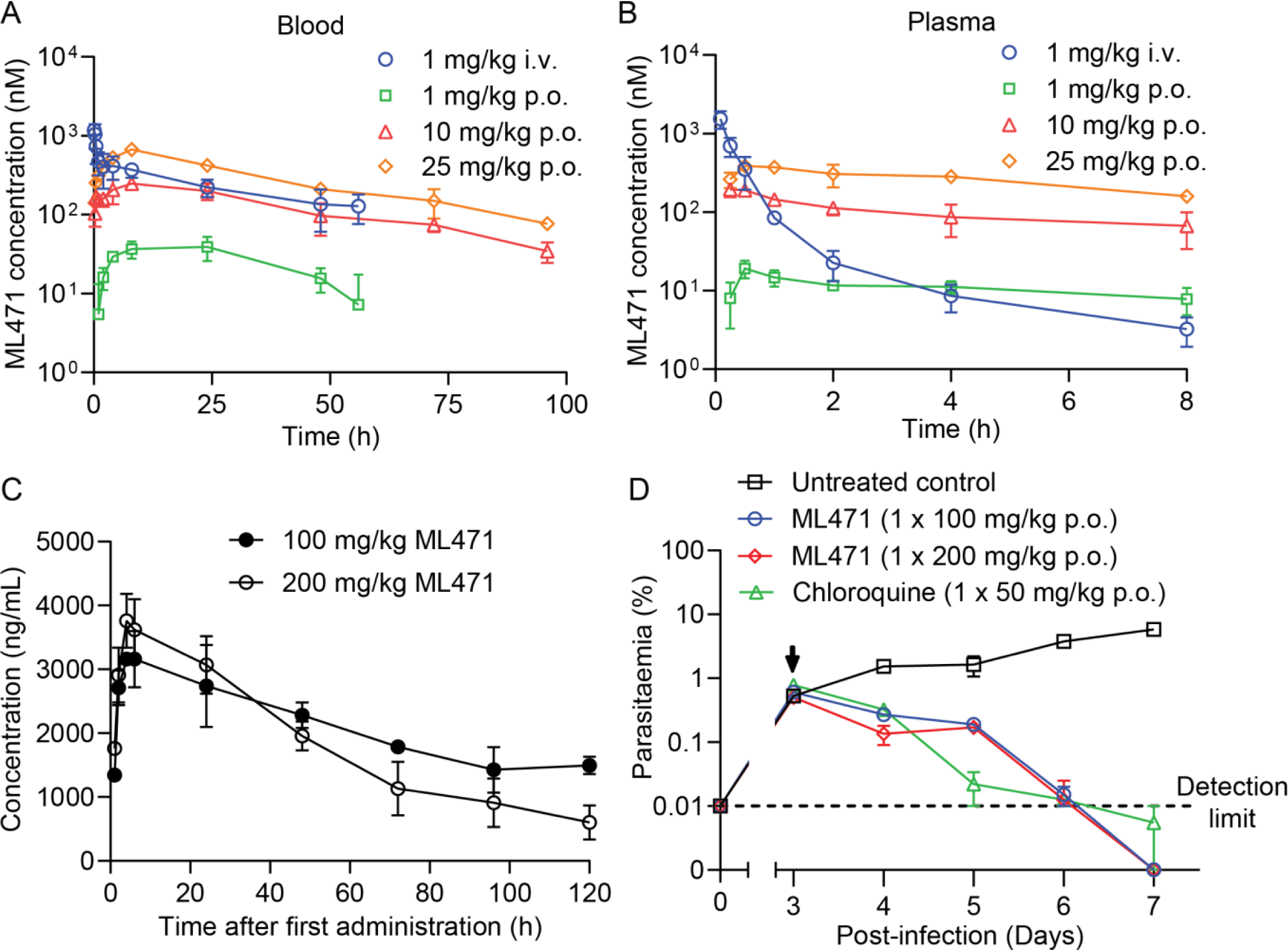
Pharmacokinetics profiles and *in vivo* efficacy of ML471. (A, B) Rat pharmacokinetics for ML471. Rats were dosed with ML471 at 1 mg/kg i.v. (blue) or 1, 10 or 25 mg/kg p.o. (green, red, orange) and plasma (A) and blood (B) samples were collected for analysis. See Supplementary Table S7 for pharmacokinetics values. (C) Pharmacokinetics profile (in blood), for SCID mice engrafted with human RBCs infected with *P. falciparum*, over the first day following treatment with ML471 at 100 or 200 mg/kg p.o.. See Supplementary Table S8 for pharmacokinetics values. (D) Therapeutic efficacy of ML471 in the SCID mouse *P. falciparum* model, dosed with ML471 at 100 or 200 mg/kg p.o. on Day 3 post-infection (arrowed). The chloroquine data are from [16].

### Efficacy of ML471 in a SCID model of P. falciparum

Single, low-dose oral efficacy is a key requirement for new antimalarial treatments to be used as part of the MMV’s Single Encounter Radical Cure and Prophylaxis (SERCAP) target product profile (TPP1) [8]. We determined the *in vivo* antimalarial efficacy of ML471 in severe combined immune deficient (SCID) mice, engrafted with human RBCs and infected with *P. falciparum* [30, 31], which is the gold standard for testing *in vivo* efficacy of malaria drug candidates. Oral dosing results in good exposure. Single doses of 100 mg/kg p.o. and 200 mg/kg p.o. resulted in AUC_0-120h_ values of 640 μM.h (250,000 h.ng/mL) and 550 μM.h (215,000 h.ng/mL), respectively (Fig. 3C; Table S8), while a dosing regimen of 4 x 50 mg/kg p.o. resulted in an area under the curve (AUC_0-24h_) of 54 μM.h (21,100 h.ng/mL), assessed at 24 h, after the first dose) (Fig. S2A; Table S8). All doses were well tolerated, and the long half-life and high exposure are encouraging as these are important properties for single dose antimalarials.

Mice were infected intravenously with 2 x 10^7^ *P. falciparum* (*Pf*3D7^0087/N9^) and ML471 was administered on day 3 post-infection. The single dose regimen (either 100 or 200 mg/kg p.o.) was sufficient to achieve reduction of 3D7 parasitemia to baseline, with a parasite clearance rate similar to that of chloroquine (CQ; 50 mg/kg p.o.) (Fig. 3D) and no evidence of toxicity. Similarly, the dosing regimen of 4 x daily doses of 50 mg/kg p.o. reduced the 3D7 parasitemia to baseline with a clearance rate similar to chloroquine (CQ; 4 x 50 mg/kg p.o.) (Supp Fig. 2B).

### ML471 and ML901 selection leads to amplification of the PfTyrRS locus

*In vitro* evolution of resistance, under a standardized protocol, has been used to assess the propensity for the development of resistance [32–34]. Here, we examined the resistance potential of both ML901 and ML471, employing a single-step selection with Dd2-B2 parasites. For ML901, the parasites were subjected to pressure at 3 x IC_50_, while for ML471, parasites were subjected to 10 x IC_50_. With both compounds, parasites were retrieved and IC_50_ shifts were observed (ranging from two- to 16-fold; Table S9). The Minimum Inoculum for Resistance values for ML901 and ML471 were estimated to be 10^7^ and 7.1 x10^5^, respectively (Table S9). These values are at or below the preferred threshold for further development, making this a parameter of concern. Use of these sulfamates in a drug combination could suppress the evolution of resistant mutants.

We performed whole-genome sequencing of parasite lines selected for resistance. Copy number variations (CNVs) were found in flasks selected with either ML901 or ML471, with amplifications always containing the *Pf*TyrRS gene located within amplicons of varying sizes (Table S10, S11). This gene was present in 2-4 copies in the amplified lines. This finding is consistent with our earlier identification of *Pf*TyrRS as the target of ML901 [16]. No SNPs were found in any of the samples, in contrast to a previous report [16]. This may be due to the slow ramp-up exposure method employed in the earlier study.

### ML471 targets P. falciparum tyrosine tRNA synthetase via a reaction hijacking mechanism

Reaction hijacking inhibition of *Pf*TyrRS is expected to lead to the formation of Tyr-ML471 adducts (Fig. 1K) in the active site. We treated *P. falciparum* infected RBCs for 2 h with 1 μM ML471 and subjected extracts to LC-MS to search for amino acid-ML471 conjugates. An LC-MS peak corresponding to Tyr-ML471 precursor ion (*m/z* 552.1871) was detected at the retention time of 3.0 min (Fig. 4A). Synthetic Tyr-ML471 was generated as a standard to confirm the peak assignment (Fig. 4A, Supplementary Fig. 3A). None of the other 19 possible amino acid conjugates were detected.

**Figure 4.**
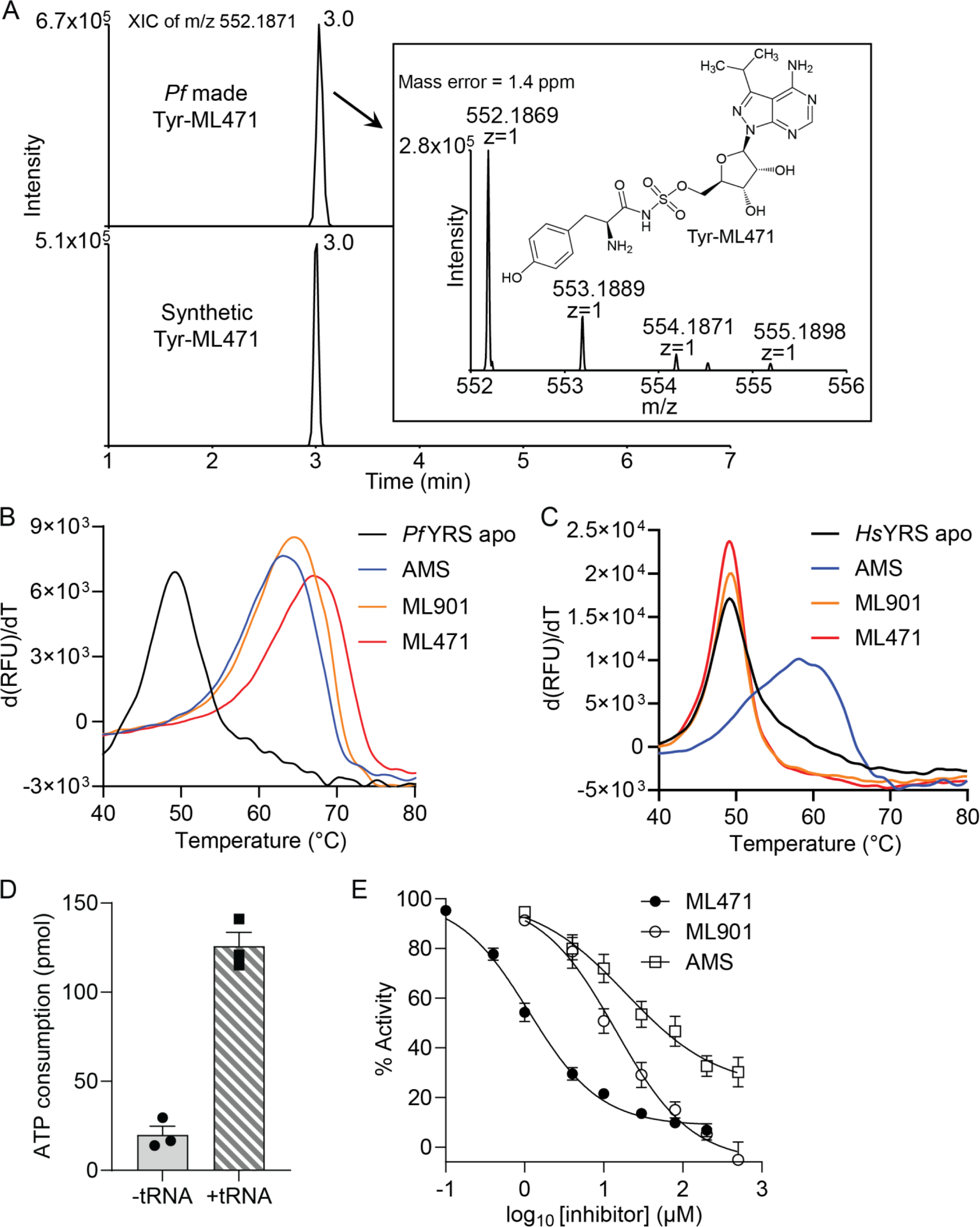
Identification of ML471 conjugates in *P. falciparum* and effects of pro-inhibitors on enzyme stability and activity. *P. falciparum*-infected RBCs were treated with 1 μM ML471 for 2 h. Extracts were subjected to LCMS and the expected mass for amino acid-ML471 conjugates searched. (**A**) The extracted ion chromatograms of the Tyr-ML471 adduct made by a *P. falciparum* culture (upper panel) and the synthetic conjugate at 0.2 μM (lower panel). The inset shows the MS analysis of the parasite-generated Tyr-ML471, and the structure of Tyr-ML471. Profiles are typical of data from 3 independent experiments. (B,C) First derivatives of melting curves for *Pf*TyrRS (B) and *Hs*TyrRS (C) (2.3 μM) in the apo form or after incubation at 37°C with ML901, ML471 or AMS, in the presence of 10 μM ATP and 20 μM tyrosine. For *Pf*TyrRS, 50 μM pro-inhibitor and 4 μM *Pf*tRNA^Tyr^ were incubated with substrates for 2 h. For *Hs*TyrRS, 200 μM pro-inhibitor and 8 mg/mL yeast tRNA were incubated with substrates for 4 h. Data are representative of three independent experiments. (D) ATP consumption by *Pf*TyrRS in the presence and absence of the cognate tRNA^Tyr^. ATP consumption in the absence of tRNA^Tyr^ derives from turnover of Tyr-AMP generated in the initial phase of the TyrRS reaction. The reaction component concentrations are: *Pf*TyrRS (25 nM), ATP (10 μM), tyrosine (200 μM), pyrophosphatase (1 unit/mL) and cognate tRNA^Tyr^ (4.8 μM), if present; and incubations were at 37°C for 1 h. Data are the average of three independent experiments and error bars correspond to SEM. (E) Effects of increasing concentrations of ML471, ML901 and AMS on ATP consumption by *Pf*TyrRS. Assay conditions are the same as in (D), with cognate tRNA^Tyr^. Data represent mean ± SEM from three or four independent experiments.

Recombinant *Pf*TyrRS was produced in *Escherichia coli* as previously described [16]. To examine the ability of *Pf*TyrRS to generate the Tyr-ML471 conjugate, the enzyme was incubated with ATP, Tyr, tRNA^Tyr^ and ML471. Following sample extraction, LC-MS analysis revealed a peak at *m/z* 552.1868 with a retention time of 3.0 min, consistent with that of the Tyr-ML471 precursor ion (Supplementary Fig. 3B). MS/MS analysis further confirmed the identity of the conjugate (Supplementary Fig. 3C).

### Recombinant P. falciparum tyrosine tRNA synthetase is thermally stabilised upon formation of the Tyr-ML471 adducts

When ML471 was incubated with *Pf*TyrRS in the presence of all other substrates (*i.e.,* Tyr, ATP and *Pf*tRNA^Tyr^), the apparent protein melting point (T_m_), measured by differential scanning fluorimetry (DSF), increased by a remarkable 18°C (Fig. 4B, Table 3). The increase in thermal stability is even greater than that induced by the Tyr-ML901 adduct (Table 3), suggesting higher affinity binding, which is consistent with the enhanced antimalarial activity of ML471 compared with ML901. Importantly, when incubated in presence of ML901 or ML471, and all substrates, recombinant *Hs*TyrRS was not stabilised (Fig. 4C, red and orange curves). This shows that the human enzyme is not susceptible to hijacking by ML471. By contrast, incubation of *Hs*TyrRS with the broad specificity compound, AMS, and all substrates, leads to substantial thermal stabilization, consistent with efficient reaction hijacking by AMS (Fig. 4C, Table 3).

**Table 3.** Activities of pyrazolopyrimidine sulfamates in selected biochemical assays. AMS = Adenosine 5’-sulfamate. n = Number of biological repeats. Data values represent mean ± SEM. The T_m_ values for *Pf*TyrRS and *Hs*TyrRS (2.3 μM) was measured in the apo form or after incubation at 37°C for 2 h with the pro-inhibitors (50 μM with *Pf*TyrRS and 200 μM with *Hs*TyrRS) in the presence of 10 μM ATP, 20 μM tyrosine, 4 μM cognate tRNA^Tyr^ (*Pf*TyrRS) or 8 mg/mL yeast tRNA (*Hs*TyrRS). K_D_ values are estimated from differential scanning fluorimetry (DSF) analysis using an irreversible protein thermal unfolding model that has been described previously [58].

| **Compound** | Inhibition of *Pf*TyrRS (Kinase Glo) | *Pf*TyrRS binding (DSF) (n = 3) |  | *Hs*TyrRS binding (DSF) (n = 3) |  |
| --- | --- | --- | --- | --- | --- |
|  | IC<sub>50</sub> (μM) | Delta T<sub>m</sub> (°C) | apparent K<sub>D</sub>* (x10<sup>-9</sup>) (M) | Delta T<sub>m</sub> (°C) | apparent K<sub>D</sub> (x10<sup>-9</sup>) (M) |
| **ML901** | 13 ± 3 (n = 3) | 15.2 ± 0.1 | 0.7 | 0.4 ± 0.1 | N/A |
| **ML471** | 1.4 ± 0.2 (n = 3) | 18.0 ± 0.1 | 0.2 | 0.06 ± 0.04 | N/A |
| **ML676** | 48 ± 16 (n = 5) | 15.69 ± 0.01 | 0.6 | N/A | N/A |
| **ML681** | 23 ± 8 (n = 3) | 16.4 ± 0.1 | 0.4 | N/A | N/A |
| **ML723** | 14 ± 5 (n = 3) | 16.2 ± 0.1 | 0.5 | N/A | N/A |
| **ML107** | 24 ± 5 (n = 3) | 13.6 ± 0.6 | 1.4 | N/A | N/A |
| **ML470** | 41 ± 23 (n = 4) | 15.7 ± 0.3 | 0.6 | N/A | N/A |
| **ML864** | 8.5 ± 4.5 (n = 4) | 17 ± 0.1 | 0.3 | N/A | N/A |
| **ML111** | 34 ± 14 (n = 3) | 15.1 ± 0.2 | 0.7 | N/A | N/A |
| **AMS** | 52 ± 16 (n = 4) | 13.6 ± 0.7 | 1.3 | 9.1 ± 0.1 | 8.0 |

### ML471 inhibits ATP consumption by PfTyrRS

Recombinant *Pf*TyrRS consumes ATP at a moderate level, even in the absence of tRNA, due to the generation and release of AMP-Tyr in the initial reaction phase (Fig. 4D). The rate of ATP consumption is increased 6-fold upon addition of tRNA^Tyr^, consistent with productive aminoacylation. ML471 inhibited ATP consumption by *Pf*TyrRS when added in the presence of *Pf*tRNA^Tyr^ (IC_50_ = 1.4 μM) much more potently than ML901 (IC_50_ = 13.4 μM) (Fig. 4E, Table 3). Indeed, ML471 is 6 to 30 times more effective than the other ML901 analogues examined (Supplementary Fig. 4, Table 3). AMS is also significantly less potent in this assay (IC_50_ = 51.7 μM) (Fig. 4E, Table 3).

### Docking of ML471 into UAE reveals differential interactions within the active site

Susceptibility to reaction hijacking depends on the ability of the pro-inhibitor to bind in the ATP-binding pocket of the relevant adenylate-forming enzyme, in a pose that is suitable for reaction with the relevant enzyme-bound product. To probe the molecular basis for the enhanced selectivity of ML471 compared with ML901, we used the Surflex-Dock molecular docking module in SybylX2 to dock ML901 and ML471 into the ATP-binding site of human UAE (6DC6) [35]. For comparison, ML901 and ML471 were docked into the binding pockets of the A- and B-chains of the *Pf*TyrRS/ Tyr-ML901 complex (7ROS) [16], noting that the two chains of the dimeric *Pf*TyrRS structure in 7ROS show differences in the position of key residues around the Tyr-ML901 ligand that are thought to relate to altered mobility of the KMSKS loop (Xie et al., 2022).

As described above, ML901 and ML471 bear, respectively, difluoromethoxy and isopropyl groups at the 7-postion of the pyrazolopyrimidine ring. Both ML901 and ML471 can be docked into the active sites of the A- and B-chains of *Pf*TyrRS, with the 7-position substituent located in a solvent accessible pocket. Fig. 5A illustrates the B-chain with docked ML901 (aqua carbons), overlaid with the pose adopted when ML901 (yellow carbons) is docked into the A-chain. In both cases the difluoromethoxy group is positioned away from His70 (red arrow). By contrast, when ML471 (aqua carbons) is docked into the B-chain and overlaid with the pose adopted when ML471 (yellow carbons) is docked into the A-chain, the isopropyl group is positioned closer to His70 (Fig. 5B), indicative of a clash.

**Figure 5.**
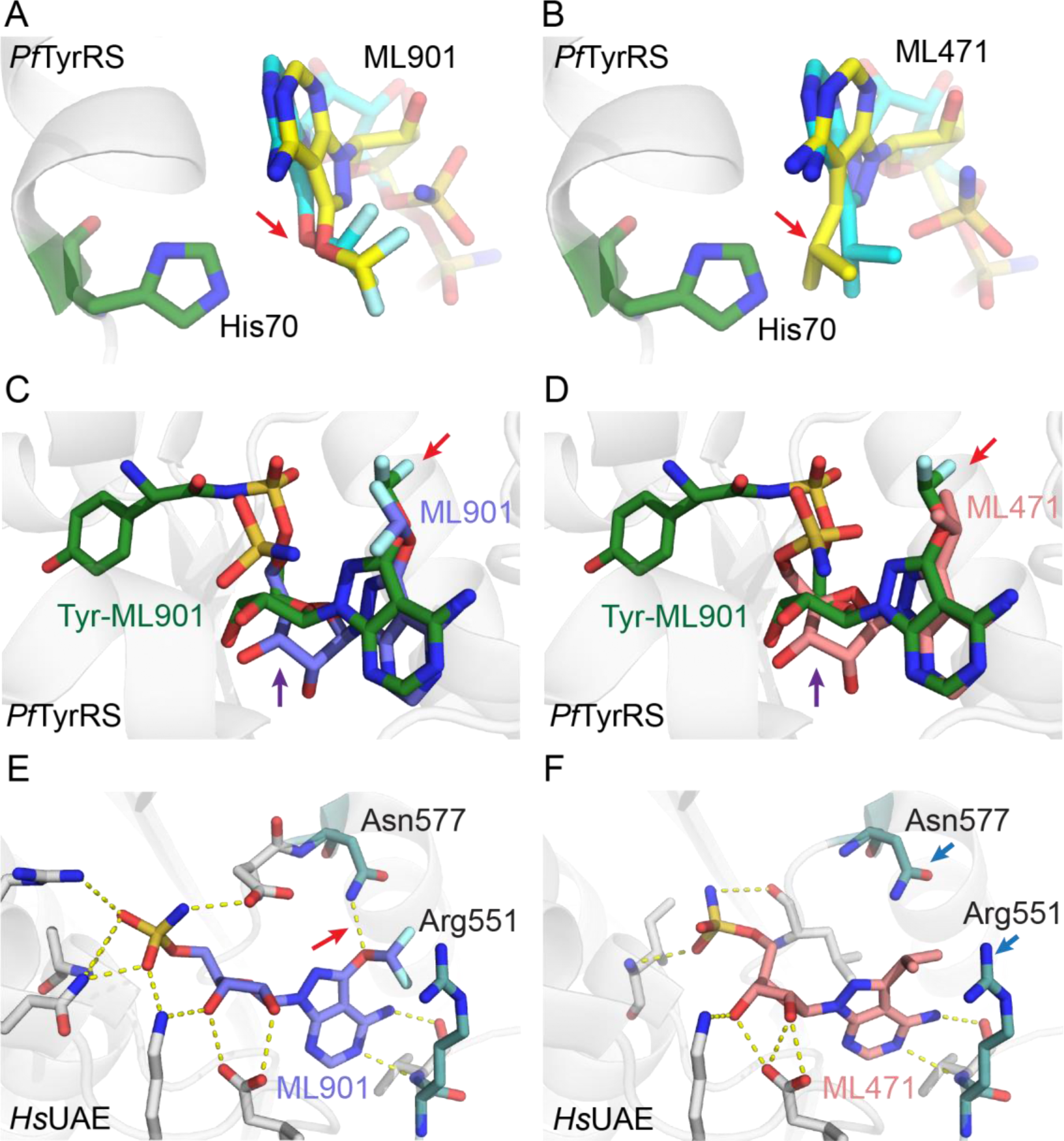
Docking of ML901 and ML471 into structures of *Pf*TyrRS and UAE provides insights into selectivity. (A) Active site of *Pf*TyrRS/Tyr-ML901 (7ROS) B-chain (His70 depicted in green) with docked ML901 (aqua carbons). The model is overlayed with ML901 (depicted with yellow carbons) with the pose adopted upon docking into the A-chain. (B) Active site of *Pf*TyrRS/Tyr-ML901 (7ROS) B-chain (His70 depicted in green) with docked ML471 (aqua carbons). The model is overlayed with ML471 (depicted with yellow backbone) with the pose adopted upon docking into the A-chain. The red arrow illustrates the different conformations adopted by the difluoromethoxy and isopropyl groups. (C, D) The structure of 7ROS B-chain with bound Tyr-ML901 is overlayed with B-chain-docked ML901 (C) and ML471 (D). The red arrows illustrate the different conformations adopted by the difluoromethoxy and isopropyl groups. The purple arrows illustrate the twisted ribose group in the Tyr-ML901 conjugate. By contrast in the docked pro-inhibitors, the rings systems are co-planar. (E,F) ML901 (E) and ML471 (F) were docked into the ATP-binding site of human UAE (6DC6). A H-bond made by ML901 with residue Arg 551 is indicated with a red arrow. Asn577 and Arg551 (blue arrows) flank the hydrophobic isopropyl group in the ML471 dock.

Overlay of the docked conformations of the two pro-inhibitors with the published *Pf*TyrRS/ Tyr-ML901 structure (7ROS) shows close alignment of the nucleoside sulfamate regions (7ROS) (Fig. 5C,D). Again, the different poses of the difluoromethoxy and isopropyl groups are evident (red arrows). Of interest, the ribose ring of Tyr-ML901 in the crystal structure is twisted relative to the docked pro-inhibitors (purple arrows). This may arise from the conjugation of the ML901 sulfamate to tyrosine, which repositions the ligand (Fig. 5C,D).

When ML901 and ML471 are docked into the ATP-binding site of human UAE, the oxygen of the ML901 difluoromethoxy group makes a H-bond interaction with Asn577 (Fig. 5E) that cannot exist for the isopropyl group in ML471 (Fig. 5F). Instead, the hydrophobic isopropyl group must pack into a polar space flanked by Asn577 and Arg551, an unfavourable interaction that is likely to reduce the affinity of ML471 for the site. The sulfamate groups are positioned slightly differently (Fig. 5E,F). However, as the chemical structure of this region of the two compounds is identical, this difference is regarded as an artefact of the docking rather than a significant alteration of the probable binding pose. Thus, differences in the interactions made by the 7-position substituent appear likely to underpin the improved selectivity of ML471.

### Crystal structure of PfTyrRS-Tyr-ML471 provides insights into the molecular basis for enhanced potency

The complex of *Pf*TyrRS with synthetic Tyr-ML471 was crystalised using previously established conditions [16] and the structure refined at 1.8 Å resolution, revealing a homodimer organization (Fig. 6A) with clear density for the Tyr-ML471 ligand. As described previously [16], *Pf*TyrRS is a Class I aaRS, characterized by a catalytic domain with Rossmann fold structure (residues 18–260) linked to a C-terminal anti-codon binding domain (residues 261–370) involved in recognition of tRNA^Tyr^. *Pf*TyrRS contains “HIGH” and “KMSKS” (_70_HIAQ_74_ and _247_KMSKS_251_ in *Pf*TyrRS) motifs that are characteristic of the catalytic domain of Class I tRNA synthetases (sub-class c).

**Figure 6.**
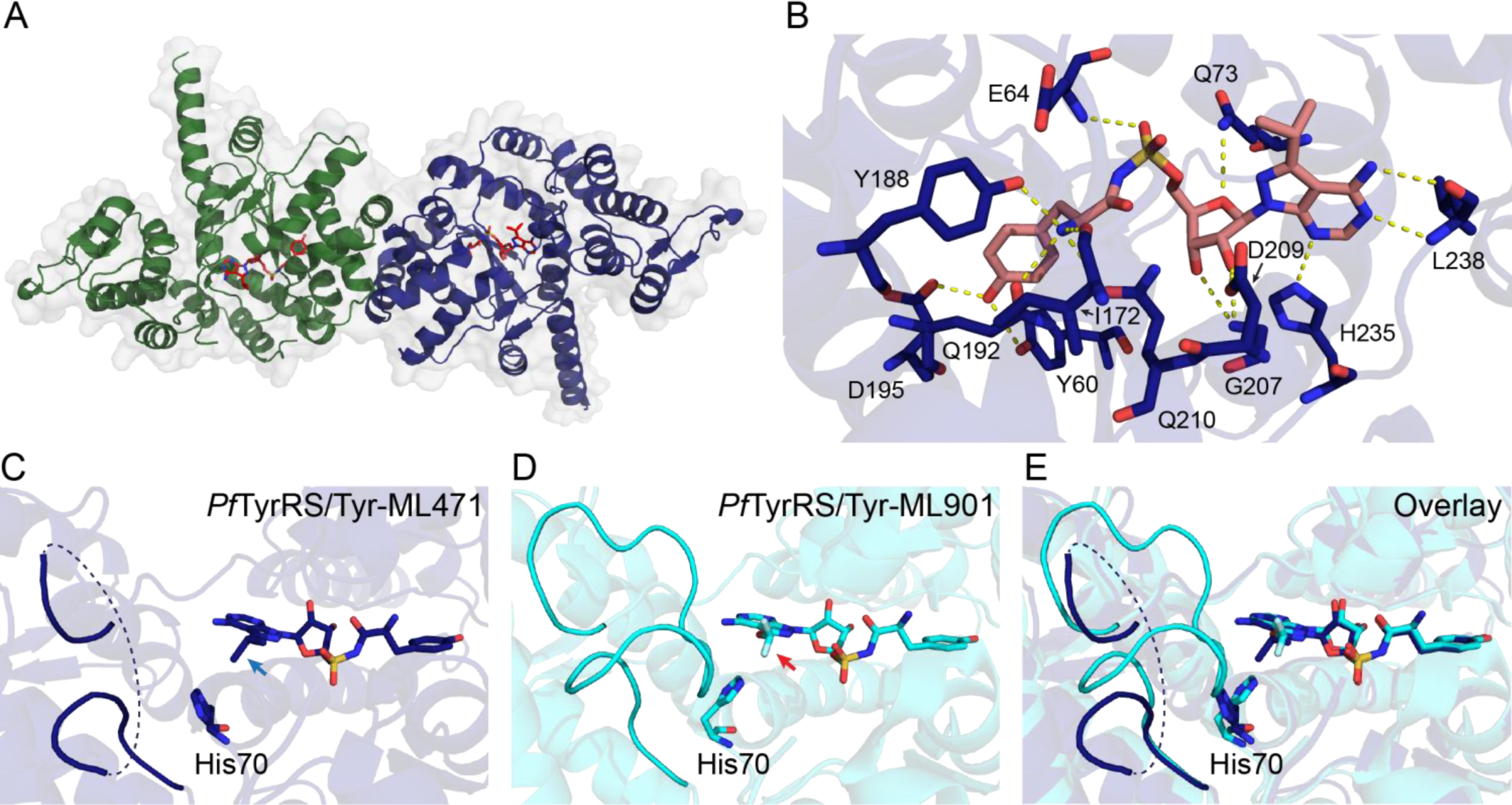
Comparison of the crystal structures of Tyr-ML471- and Tyr-ML901-bound *Pf*TyrRS reveals differential mobility of the “KMSKS” loop. (A) Crystal structure of the dimeric *Pf*TyrRS/Tyr-ML471 complex showing chain A (green), chain B (blue), and bound Tyr-ML471 (red, stick representation). (B) Architecture of the B-chain of *Pf*TyrRS with bound Tyr-ML471, showing direct interactions with active site residues. (C) B-chain of Tyr-ML471-bound *Pf*TyrRS showing the poses adopted by the ML471 isopropyl group (blue arrow) and His70 (H70), which are incompatible with a structured KMSKS loop. (D) B-chain of Tyr-ML901-bound *Pf*TyrRS (7ROS). The conformation of the ML901 difluoromethoxy group (red arrow) allows His70 to interact with Met248 of the KMSKS loop, leading to stabilisation. (E) Overlay of the B-chains of Tyr-ML471- and Tyr-ML901-bound *Pf*TyrRS.

Tyr-ML471 makes many interactions with active site residues, involving the pyrazolopyrimidine amine, ribose hydroxyls, sulfamate, and tyrosine (Fig. 6B; Supp Fig. 5A,B). The 7-position isopropyl group is oriented away from the binding pocket and is partially solvent exposed, consistent with the docking study. The isopropyl group of ML471 is oriented differently in the A- and B-chains and the adjacent His70 (of _70_HI_AQ73_) adopts different side chain rotamers in each chain (Fig. 6C; Supp Fig. 5C). The majority of the KMSKS loop was poorly defined in the electron density in both chains, suggesting flexibility.

Comparison of the B chains of the Tyr-ML901-bound and Tyr-ML471-bound *Pf*TyrRS structures highlights a difference in the KMSKS loop organisation. The difluoromethoxy group of ML901 is oriented away from His70 (Fig. 6D, red arrow), allowing His70 to adopt a configuration that makes close contact with Met248 in the _247_KMSKS_251_ loop, thereby stabilising the loop (Fig. 6D,E; aqua backbone). By contrast the isopropyl group of ML471 in chain B is oriented with one methyl group towards His70 (Fig. 6C, blue arrow), and the His70 side chain adopts a rotamer that is incompatible with the Met248 interaction observed for Tyr-ML901. Thus, for chain B, Tyr-ML471 binding appears to be associated with loop destabilisation while Tyr-ML901 binding is associated with loop stabilisation. Interestingly, in the A-chain, His70 adopts a similar pose that does not support interaction with Met248 in both Tyr-ML471- and Tyr-ML901-bound structures, leading to destabilisation of the loop (Supp Fig. 5C-E).

Movement of the KMSKS loop is required for access to the active site. For example, in tRNA^Trp^-bound human TrpRS, the equivalent KMSAS loop adopts a semi-open conformation, that is intermediate between the open conformation observed in the unliganded enzyme and the closed conformation observed in the Trp-AMP complex [36]. The pose adopted by the 7-position isopropyl group of ML471 appears to re-position His70, leading to increased flexibility of the KMSKS loop. This may, in turn, enhance the binding or re-binding of the Tyr-tRNA product, positioning the Tyr-tRNA carbonyl carbon for attack by the sulfamate nitrogen of ML471. This may underpin the higher potency of ML471 as a hijacking inhibitor of recombinant *Pf*TyrRS and as an inhibitor of the growth of *P. falciparum*.

In conclusion, we have identified ML471 as a pyrazolopyrimidine sulfamate with improved potency and selectivity compared with ML901. The improved potency derives from improved efficiency of reaction hijacking inhibition of *Pf*TyrRS and/ or tighter binding of the Tyr-ML471 adduct, as indicated by enhanced thermal stability. The enhanced potency of ML471 is associated with repositioning of His70 in the active site of chain B when Tyr-ML471 is bound. The enhanced cellular selectivity of ML471 may be due to its decreased activity against human UAE, which is associated with a lack of interaction between the 7-position substituent of ML471 and active site residues. ML471 exhibits a rapid mode of action and a long *in vivo* half-life underpinning its single dose efficacy in a mouse model of *P. falciparum* malaria. With further improvement of the oral bioavailability, ML471 represents a very interesting compound for prophylaxis, treatment and blocking of transmission of deadly malaria infections.

## Materials and methods

### Inhibition of growth of P. falciparum cultures

Routine analyses of antimalarial activity against *P. falciparum* 3D7 was tested by TCGLS (Kolkata, India) using the lactate dehydrogenase (*Pf*LDH) growth inhibition assay as previously described [37]. For assays readout, 70 μL of freshly prepared reaction mix containing 100 mM Tris-HCl pH 8, 143 mM sodium L-lactate, 143 μM 3-acetyl pyridine adenine dinucleotide (APAD), 179 μM Nitro Blue tetrazolium chloride (NBT), diaphorase (2.8U/mL) and 0.7% Tween 20 was added into each well of the assay plate. Plates were shaken to ensure mixing and were placed in the dark at 21⁰C for 20 min. Data were normalized to the percentage of growth inhibition with respect to positive (0.2% DMSO, 0% inhibition) and negative (mixture of 100 μM chloroquine and 100 μM atovaquone, 100% inhibition) controls. *P. falciparum* strain (3D7) was obtained from BEI Resources.

### Analysis of inhibition of growth and viability of P. falciparum cultures followed pulsed compound exposure

Assessment of the killing activity of ML901 in pulsed exposure assays was performed using a modification of a previously described procedure [38]. Briefly, cultures of Cam3.II^rev^ [39] trophozoites (1.5% final hematocrit; 1.4% final parasitemia) were pre-synchronized to a 5-h window at trophozoite or schizont stage [40]. Compounds were serially diluted in complete medium in v-bottomed microplates. Parasites were exposed to ML901 for 3 h, 6 h, 9 h or 24 h before washing 5 times with 200 µl of complete medium, then returned to culture conditions. For the no wash samples, ML901 was left in the culture until the assay point. Growth inhibition was assessed in the second cycle by labelling with the DNA-binding dye, SYBR Green I. Quantification of total DNA level reflects cytostatic effects as well as cytocidal effects [41]. Old media (140 µl) was firstly replaced with fresh media, followed by the addition of lysis buffer (20 mM Tris, pH 7.5, 5 mM EDTA, 0.008% w/v saponin, 0.08% v/v Triton X-100) containing SYBR Green I. Plates were incubated at room temperature for 2 h and fluorescence readings were taken using a microplate reader (BMG LABTECH). Unwashed samples containing compounds at >10 times the IC_50_ values were used as background controls.

### Activity against HepG2 cell cultures

The HepG2 (Human Caucasian hepatocyte carcinoma) cell line was procured from ATCC (American Type Culture Collection, Manassas, USA; HB-8065) and viability assessed using CellTiter-Glo luminescent cell viability assay reagents (Promega). For the assay, 2,000 cells/well were plated in 384-well plates 24 h prior to the experiment and incubated at 37°C in a CO_2_ incubator. The medium was removed; and cells were treated with fresh medium containing either vehicle (0.5% DMSO) or serially diluted compounds or doxorubicin (1.3 nM to 25 μM) in a final volume of 50 μL/well and further incubated for 72 h at 37°C in a CO_2_ incubator. In the positive control wells (100% inhibition), cells were treated with 5 μL of 1% Triton X-100. Following incubation, 25 μL of medium was discarded and 25 μl of CellTiter-Glo reagent was added to each well and the plate was kept on a plate shaker for 15 min at 25°C with shaking at 300 rpm. Luminescence signals were measured in a PHERAstar FSX reader (BMG LABTECH).

### E1-E2 Transthiolation Assays

An Homogeneous Time-Resolved Fluorescence (HTRF) enzyme assay was employed to evaluate compound activity against ATG7 as previously described [12]. In this assay, a Flag-tagged ATG8 homolog (GABA type A receptor-associated protein; GABARAP) is activated by ATG7 and then transthiolated to a GST-tagged E2 (ATG3). The product of the enzyme reaction, *Flag*-GABARAP-ATG3-*GST*, is quantified by measuring FRET between Europium-Cryptate labelled monoclonal anti-Flag M2 (FLAG M2-Eu cryptate; Revvity, Cat# 61FG2KLB) and goat polyclonal antibody against mouse IgG conjugated to allophycocyanin (APC) (Anti-GST IgG conjugated to SureLight™-Allophycocyanin) (Revvity, Cat# AD0059G)). The activation and transthiolation of ubiquitin by UAE, activation and transthiolation of NEDD8 by NAE and activation and transthiolation of SUMO by SAE were all assayed in a similar fashion with appropriately tagged ubiquitin-like proteins and E2 conjugating enzymes as described [9, 25].

### Activity against panels of ex vivo field isolates of P. falciparum and P. vivax

Compounds were assayed on ten *P. vivax* isolates and seven *P. falciparum* Brazilian isolates collected from mono-infected patients, who had signed a written informed consent form to participate in the study, using previously described methods [42]. The initial parasitemia ranged from 2,100-8,000 parasites/μL for *P. vivax* and 3,500-9,000 parasites/μL for *P. falciparum* isolates. Artesunate and chloroquine were assayed in parallel as standard compounds. The analyses included only the isolates that were incubated for ≥ 40 h with the compounds.

### Parasite Reduction Rate (PRR) estimation

PRR was assessed using a standardized method [27]. *P. falciparum* (strain 3D7A, MRA-151), contributed by David Walliker, was obtained through BEI Resources, NIAID, NIH. Cultures of parasites (∼90% ring stage) were treated with compounds for 120 h, with daily renewal. Samples of parasites were taken at 0, 24, 48, 72, 96 and 120 h. Compounds were washed out and four independent 3-fold serial dilutions were established in 96-well plates, with fresh RBCs and culture medium. After 18 days, and again at 22 days, samples were taken to examine growth using SYBR Green I in an EnVision Multilabel Plate Reader (Perkin Elmer) and analysed using Excel and Grafit 5.0 software. The human biological samples were sourced ethically, and their research use was in accord with the terms of the informed consents under an IRB/EC approved protocol.

### Minimum inoculum of resistance

Minimum Inoculum of Resistance (MIR) studies were conducted for ML901 and ML471 using a modified “Gate keeper assay” [34]. The in-house IC_50_ for ML901 2.6 ± 0.05 nM (mean ± SEM; N,n = 2,2), while for ML471 the mean IC_50_ and IC_90_ values were determined as 1.45 and 1.99 nM, respectively. For ML901, the parasites (starting parasite inoculums of 1×10^7 or 1×10^8 in triplicate) were subjected to pressure at 3 x IC_50_, with recrudescence on days 12-14. For ML471, parasites were subjected to 10 x IC_50_. Wells were monitored daily by smear during the first seven days to ensure parasite clearance, during which media was changed daily. Thereafter, cultures were screened three times weekly by flow cytometry and smearing, and the selection maintained a consistent drug pressure over 60 days. In both cases, IC_50_ shifts were observed (ranging from two- to 16-fold). Whole-genome sequencing analysis identified CNVs in chromosome 8 segments, always containing the *Pf*TyrRS locus, consistent with our earlier evidence of this as the target of ML901 [16]. Single nucleotide polymorphism (SNP) filtering was lowered to 0.5 allelic balance (AB) but yielded no point mutations in the core genome.

### Gametocyte killing assays

Gametocytogenesis was induced on a tightly synchronised (>97% rings) asexual parasite culture (Pf3D7-pfs16-CBG99 (kind gift from Pietro Alano), 0.5% parasitemia and 6% hematocrit) with a combination of nutrient starvation and a decrease in hematocrit, as previously described [43]. For immature gametocytes (>90% stage II/III), cultures were exposed to 50 mM N-acetyl glucosamine (NAG) on days 1–4 to eliminate residual asexual parasites and harvested at days 5–6. For mature (>95% stage V) gametocytes, NAG treatment occurred from days 3–7 and harvested at day 13. Stage II/III and V gametocyte cultures (2 % gametocytaemia 1.5 % haematocrit, 150 μL/well) were exposed to compounds and incubated at 37°C for 48 h under hypoxic conditions [44], after which luciferase activity was determined with a non-lysing D-luciferin substrate (1mM in 0.1M citrate buffer, pH 5.5, 100 µL) and bioluminescence was detected with a 2 s integration time with a GloMax^®^-Multi Detection System with Instinct^®^ software.

### Activity against P. falciparum male and female gamete formation

Inhibition of the viability of stage V male and female gametocytes was assessed in the *P. falciparum* Dual Gamete Formation Assay (*Pf*DGFA) as described previously [45]. Briefly, mature stage V NF54 strain gametocytes were incubated with test molecules for 48 h in 384-well plates at 37°C. Gametogenesis was then triggered by cooling the plates to room temperature and addition of xanthurenic acid-containing activating medium (also containing anti-*Pf*s25 antibody (Mab 4B7) conjugated to a Cy3 fluorophore). Twenty minutes after activation, exflagellation was imaged in all wells of the plate using a x4 objective and automated brightfield microscopy. The plate was then incubated at 20°C for a further 24 h in the dark. Thereafter, female gamete formation was quantified by automated identification of *Pf*s25-positive cells. Automated counts were transformed to percent inhibition values with reference to positive 100 nM Cabamiquine (DDD498) and negative (DMSO) controls. Data represent the means of multiple independent biological repeats.

### Activity against liver stage P. falciparum

Activity against liver stage parasites was performed using a modification of published procedures [46, 47]. Briefly, cryopreserved human primary hepatocytes (NF175: H1500.H15B+ Lot No. HC0-6, TebuBio or NF135: F00995-P Lot No. IRZ, BioIVT) were thawed and seeded at 18,000 cells per well in collagen-coated 384-well microtiter plates in medium containing 10% heat inactivated fetal bovine serum (hiFBS). Cells were cultured at 37°C in 5% CO_2_. For NF175, medium was replaced by fresh medium containing 10% hiFBS, 5 h post plating. For NF135, medium was replaced by medium containing 0.2% BSA, 24 h post plating. 48 h post plating, salivary glands from *Plasmodium*-NF175 or NF135-infected *Anopheles stephensi* mosquitoes were dissected, added to the hepatocytes (10,000 per well/ NF175; or 12,500 per well/ NF135) and allowed to infect for 3 hours. Sporozoites were then aspirated, and compounds diluted in medium containing 10% hiFBS or 0.2% BSA, were added to the hepatocytes. Medium containing 10% hiFBS or 0.2% BSA and compounds was refreshed daily for four days. Hepatocytes were fixed with ice-cold methanol and samples were blocked with 10% hiFBS in PBS. Schizonts were stained with rabbit anti-HSP70 in 10% hiFBS for 1-2 h followed by incubation with a mixture of secondary goat anti-rabbit AlexaFluor 594 antibody and 4′,6-diamidine-2′-phenylindole dihydrochloride (DAPI) in 10% hiFBS for 30 min. Samples were washed with PBS containing 0.05% Tween 20 between different steps. Cells were imaged on a PicoExpress high content imager and images were analysed automatically using CellReporterXpress software. Data were analysed by logistic regression using a four-parameter (Hill equation) model and a least-squares method to find the best fit.

### Rat pharmacokinetics (PK) analyses

Sprague-Dawley rats (11 weeks old) were sourced from Hilltop Lab Animals, Inc (Scottdale, Pennsylvania, USA). ML901 was formulated in ethanol: dimethyl acetamide (DMAc): PEG400: H_2_O at 1:1:4:4 (v/v) and 10% captisol in 50 mM citrate (pH 3.3) for i.v. (1 mg/kg) and p.o. (10 mg/kg) administration to male Sprague-Dawley rats (n = 3 per route of administration). Blood was collected from a jugular cannula at 0.083, 0.25, 0.5, 1, 2, 4, 8 and 24 h post i.v. dosing, and at the same times (except the 0.083-h sample) following oral administration. A portion of the blood samples were processed into plasma. Samples were precipitated with 0.5% formic acid in methanol and the supernatants were analysed by positive ion electrospray LC-MS for the administered compound. Non-compartmental pharmacokinetic parameters were calculated from individual concentration *vs* time profiles using Phoenix 64 (WinNonlin) Version 8.1 Certara, Princeton NJ.

### Permeability analysis

Permeability studies were performed as described previously [48]. In brief, Caco-2 cells were cultured for 21-25 days to differentiate them into enterocyte-like cells. The transepithelial electrical resistance (TEER) was measured to ensure tight junction formation and cells with TEER value more than 250 ohms.cm^2^ were used in the study. On the day of the transport study, cells were washed with warm HBSS buffer and equilibrated with buffer for 60 min. ML471 was added at a concentration of 5 µM (containing 50 µM Lucifer Yellow) into a 24 Transwell cell plate (apical 210 μL and basal 1000 μL) and buffer was added in the receiver side. Cells were incubated at 37°C for 60 min and 120 μL aliquots were taken from the receiver side after 30 and 60 min. Samples were mixed with 100 nM carbutamide in acetonitrile (ACN) containing 0.1% formic acid (internal standard). Samples were centrifuged at 2,056 x g for 10 min and the supernatant was collected and analyzed for quantification of the test article by LC-MS [48].

### P. falciparum humanized NOD-scid IL2Rnull mouse model

The model using *P. falciparum Pf*3D7^0087/N9^ in NODscidIL2Rγ^null^ mice engrafted with human RBCs was adapted from a previously described procedure [49]. Female NODscidIL2Rγ^null^ mice were purchased from Charles River (Germany). Briefly, two engrafted mice/dosing group (females, 20 - 22 g) were infected intravenously with 2 x 10^7^ *P. falciparum* (*Pf*3D7^0087/N9^) on day 0. The antimalarial efficacy was assessed following administration (p.o.) of 100 or 200 mg/kg of compound or of 4 daily doses of 50 mg/kg, initiated on day 3 post-infection. The effect on blood parasitemia was measured by microscopic analysis of Giemsa-stained blood smears (on days 3, 4, 5, 6 and 7 post-infection). Mice were euthanized on day 7.

### Plasma exposure in the infected mouse model

Compound was administered orally to two mice at 25 mg/kg on days 3, 4, 5, 6 after infection. On day 3, blood samples (20 μL) were obtained at time points up to 24 h after the first administration. Protein was precipitated with acetonitrile and the remaining compound was assessed by LC-MS/MS in the selected reaction monitoring mode using HESI ionization in positive ion mode.

### Preparation of P. falciparum TyrRS

*Pf*TyrRS was expressed and purified as previously reported [16]. Briefly, the vectors were transformed into *E. coli* BL21(DE3) cells and induced for 3 h at 37°C with 0.1-0.5 mM IPTG. Cell pellets were resuspended in the lysis buffer containing 50 mM Tris-HCl, pH 8, 500 mM NaCl, 50 mM imidazole, 1 mM TCEP, 1 mg/mL lysozyme and 1x protease inhibitor cocktail (Roche). Cells were lysed by sonication (Microtip, QSonica) and the lysate was clarified by centrifugation. The supernatant was applied to a HisTrap HP column (GE Healthcare), washed, and eluted using a 0-500 mM imidazole gradient. The eluted His-*Pf*TyrRS was dialyzed overnight with the addition of His-tagged TEV protease (L56V/S135G/S219V triple-mutant). Cleaved His tags and TEV protease were removed by running the dialyzed protein through a HisTrap HP column. *Pf*TyrRS was further purified by gel filtration using a HiLoad 16/600 Superdex 200 column (GE Healthcare).

### In vitro transcription/ translation of PftRNA^Tyr^

A T7 RNA polymerase promoter sequence was added to the 5’ end of the DNA sequence of *PftRNA^Tyr^*. This DNA template and its complementary strand were custom-synthesised by Sigma-Aldrich. Two oligonucleotides were annealed at 95°C for 5 min and the double-stranded DNA template was used for *in vitro* transcription. The transcription reaction was incubated at 37°C overnight. The reaction mixture consists of template DNA, T7 RNA polymerase and NTP mix as per manufacturer’s instructions (HiScribe™ T7 Quick High Yield RNA Synthesis Kit, NEB). On the following day, the reaction mixture was treated with DNase I at 37°C for 15 min. tRNA was purified using Phenol: Chloroform: Isoamyl alcohol (25:24:1, v/v), followed by 1M LiCl precipitation and isopropanol precipitation. Purified *PftRNA^Tyr^* was subjected to NAP-25 desalting columns (Cytiva) to remove floating NTPs. The obtained solutions were concentrated with isopropanol precipitation and the *PftRNA^Tyr^* pellets were dissolved in DEPC-treated water.

### ATP consumption assays

Consumption of ATP was measured using a luciferase-based assay as per the manufacturer’s instructions (Kinase-Glo Luminescent Kinase Assay, Promega). Reactions were conducted in 50 mM Tris-HCl pH 7.6, 50 mM KCl, 25 mM MgCl_2_, 0.1 mg/mL BSA, 1 mM DTT, with 200 μM L-tyrosine, 48 μM *Pf*tRNA^Tyr^, 10 μM ATP, 25 nM *Pf*TyrRS and 1 unit/mL inorganic pyrophosphatase (yeast) in the presence or absence of 0 – 200 μM of inhibitors. Reactions were incubated at 37°C for 1 h.

### Identification of amino acid-ML471 conjugates generated by ML471-treated PfTyrRS and P. falciparum cell culture

*In vitro* recombinant *Pf*TyrRS reactions were set up with 2 μM enzyme, 20 μM tyrosine, 10 μM ATP, 10 μM ML471 and 4 μM *Pf*tRNA^Tyr^. The reaction buffer consisted of 25 mM Tris, pH 8, 150 mM NaCl, 5 mM MgCl_2_ and 1 mM TCEP. The mixture was incubated at 37°C for 1 h. An equal volume of 8 M urea was added to the mixture after the incubation, followed by trifluoroacetic acid to a final level of 1%. The sample was centrifuged at 15,000 g for 10 min and the supernatant was used for mass spectrometry analysis.

For the identification of adducts in parasite culture, aliquots of late trophozoite stage *P. falciparum* (3D7 strain) culture were exposed to 1 μM ML471 for 2 h. Following treatment, parasite-infected RBCs were lysed with 0.1% saponin and the pellet was washed 3 times with ice-cold PBS. *P. falciparum* cell pellets were resuspended in one volume of water, followed by the addition of five volumes of cold chloroform-methanol (2:1 [vol/vol]) solution. Samples were incubated on ice for 5 min, subjected to vortex mixing for 1 min and centrifuged at 13,500 x g for 10 min at 4°C to form 2 phases. The top aqueous layer was transferred to a new tube and subjected to LCMS analysis.

### High-performance liquid chromatography (HPLC) and mass spectrometric (MS) analyses

Samples were analysed by reversed-phase ultra-high performance liquid chromatography (UHPLC) coupled to tandem mass spectrometry (MS/MS) (Q Exactive, ThermoFisher Scientific). Samples (5 μL) were injected onto a Dionex Ultimate 3000 UHPLC system (ThermoFisher Scientific) and analytical separation was performed with a RRHD Eclipse Plus C8 column (2.1 × 100 mm, 1.8 μm; Agilent Technologies). The system was run at a flow rate of 300 μL/min using a binary gradient solvent system consisting of 0.1% formic acid in water (solvent A) and 0.1% formic acid in acetonitrile (solvent B). The gradient profile was as follows: 0–4.5 min, 3-40% B; 4.5–4.6 min, 40–95% B; 4.6–5.5 min, 95% B; 5.5–5.8 min, 95–3% B and 5.8–8 min, 3% B. Full scan MS acquisition was performed in polarity switching mode, with the following settings: resolution 35,000, 900 AGC target 1 × 10^6^, *m/z* range 85–1275, sheath gas 50, auxiliary gas 20, sweep gas 2, probe temperature 120°C, capillary temperature 300°C and S-Lens RF level was set to 50. The spray voltage was set at 3.5 kV for positive and negative ionization modes. All data shown for the Tyr-ML471 adduct were collected using positive mode ionisation.

### Differential scanning fluorimetry (DSF)

The effect on ML471 and analogues on the thermal stability of *Pf*TyrRS was assayed as previously described [16]. Briefly, *Pf*TyrRS (2.3 μM) was incubated with 10 μM ATP, 20 μM L-tyrosine, 4 μM *Pf*tRNA^Tyr^ and various inhibitors at 37°C for 2 h. SYPRO Orange (Sigma-Aldrich; 5,000X concentrate in DMSO) was added to the reaction mixture at a final concentration of 5X. 25 μL of the sample was added into each well of a 96-well qPCR plate (Applied Biosystems). The plate was sealed and analysed using StepOnePlus Real-Time PCR system (Applied Biosystems). The samples were heated from 20°C to 90°C with a 1% continuous gradient. The thermal unfolding curve was plotted as the first derivative curve of the raw fluorescence values. The melting temperature (T_m_), defined as the peak of the first derivative curve, was used to assess the thermal stability of protein-ligand complexes.

### Crystallography

Recombinant *Pf*TyrRS in complex with synthetic Tyr-ML471 was crystallised using the sitting drop vapour diffusion technique at 20°C. Crystals were formed in 2.25 M sodium malonate, pH 6. Drops contained 1.5 μL of protein-ligand solution (10 mg/mL *Pf*TyrRS in Tris-HCl (25 mM, pH 8)), 100 mM NaCl, 10 mM MgCl_2_, 1 mM TCEP, 500 μM Tyr-ML471 synthetic ligand) and 1.5 μL of crystallant solution (2.25 M sodium malonate, pH 6).

Crystals were flash-cooled in liquid nitrogen directly from the crystallisation drop, and X-ray diffraction data were collected at the MX2 beamline of the Australian Synchrotron [50]. Diffraction data were indexed and integrated using XDS and analysed using POINTLESS, prior to merging by AIMLESS [51] from the CCP4 software suite [52]. Initial phase estimates for *Pf*TyrRS in complex with Tyr-ML471 were obtained by molecular replacement in PHASER [53] using modified crystal structure coordinates of *Pf*TyrRS/ Tyr-ML901 as the search model (PDB ID: 7ROS, Xie 2022). Automated structure refinement using phenix.refine [54] was followed iteratively by manual model building in COOT [55]. Structure refinement was performed using non-crystallographic torsion restraints and translation/libration screw (TLS) refinement with each chain comprising a single TLS group. Restraints for Tyr-ML471 were generated using phenix.elbow [56]. Final data collection and refinement statistics are shown in Supplementary Table 12.

### Docking

Docking was carried out with the Surflex-Dock molecular docking module in SybylX2.1 (Certara, NJ, USA). Docks were performed both with and without protein flexibility. Docking poses with the best Surflex scores were inspected in SybylX2.1 and figures generated with PyMOL.

### Chemistry

The syntheses of the compounds from the series have been reported previously [12, 22]. MMV designations for the compounds are listed in Table 1. Methods used for resynthesis of ML471; and for synthesis of Tyr-ML471 are presented in the Supplementary Material. Adenosine 5’-sulfamate (AMS) [57] and was kindly provided by Dr Derek Tan, Memorial Sloan Kettering Cancer Center.

## Ethics statement

Human biological samples were sourced ethically; and their research use was in accord with the terms of the informed consent. Animal studies were ethically reviewed by the Institutional Animal Care and Use Committee at GSK or by the ethical review process at the institution where the work was performed and carried out in accordance with relevant countries’ directives, European Directive 2010/63/EU, and institution’s and GSK’s Policy on Care, Welfare and Treatment of Animals. Parasitology work and volunteer human blood donation (from healthy adult consenting volunteers) at the University of Pretoria is covered under ethical approval from the Research Ethics Committee from Health Sciences (506/2018) and Natural and Agricultural Sciences (180000094). Studies of *P. vivax* isolates and *P. falciparum* Brazilian isolates were approved by the Ethics Committee of the Tropical Medicine Research Center - CEPEM (CAAE 61442416.7.0000.0011).

## Acknowledgements

We thank the following colleagues for technical contributions: Liver Stage Assay: Marloes de Bruijni and Rob Henderson, TropIQ, Netherlands; Caco2 Assays: Bei-Ching Chuang, Takeda Pharmaceuticals, USA; SCID mouse assay: Ursula Lehmann, Swiss Tropical and Public Health Institute, Switzerland, Christoph Siethoff, Swiss BioQuant; 3D7 parasite assays: TCG LifeSciences, Kolkata, India; Assay coordination: Delphine Baud and Anna Adam, Medicines for Malaria Venture, Switzerland; Mass Spectrometry: Shuai Nie and Nick Williamson, Melbourne Mass Spectrometry and Proteomics Facility; Crystallisation: Roxanne Smith, Bio21-WEHI Crystallisation facility; Protein Purification: Yee-Foong Mok, Melbourne Protein Facility, Bio21 Institute. We thank Hirotake Mizutani, Takeda Pharmaceuticals and Winnie Ye, University of Melbourne, for technical help. Thanks also extended to Heekuk Park, Sachel Mok and Anne-Catrin Uhlemann for whole-genome sequencing and analysis at the Columbia University Irving Medical Center. We thank Dr Derek Tan, Memorial Sloan Kettering Cancer Center, for supplying adenosine 5’-sulfamate. We thank the Australian Red Cross for supply of blood products. GSK acknowledges the Centro de Hemoterapia y Donación de Valladolid, Castilla y León, and the Centro de Transfusiones de la Comunidad de Madrid for the supply of blood samples. This research was partly undertaken at the Australian Synchrotron, part of the Australian Nuclear Science and Technology Organization, and made use of the ACRF Detector on the MX2 beamline. We thank the beamline staff for their assistance.

## Data Availability

Additional data are available in Supplementary Information. Source data are provided. The following structures have been deposited in the PDB: *Pf*TyrRS/Tyr-ML471 - PDB ID 9CLL.

## Author Contributions

Conceptualisation: S.C.X., C-W.T., C.J.M., L.M., S.W., C.D., F.J.G, C.H., D.A.F., L.R.D., S.L.B., A.E.G., S.L., M.D.W.G., L.T.; Investigation: S.C.X., C-W.T., C.J.M., L.M., S.H., S.W., Y.D., Y.H., C.D., R.G., D.E., E dl C., I.D., T.Y., A.Y.B., J.S., K.A.S., B.C., Y.K., M.S.S., T.R., M.F., M.D., J.B., K.M.J.K., R.vdL., A.C.C.A., D.B.P.; Analysis: S.C.X., C-W.T., C.J.M., L.M., S.H., S.W., Y.D., Y.H., C.D., R.G., D.E., E.dl C., I.D., T.Y., A.Y.B., J.S., K.A.S., B.C., Y.K., M.S.S., T.R., L-M.B., M.F., M.D., J.B., K.M.J.K., R.vdL., A.C.C.A., D.B.P., R.V.C.G., D.A.F., L.R.D., S.L.B., A.E.G., S.L., M.D.W.G., L.T.; Funding acquisition: S.C.X., S.W., F.J.G, C.H., L-M.B., R.V.C.G,D.J.C., D.A.F., L.R.D., S.L.B., A.E.G., S.L., M.D.W.G., L.T.; Writing: S.C.X., C-W.T., C.J.M., L.M., S.W., M.S.S., C.H., D.A.F., L.R.D., S.L.B., A.E.G., S.L., M.D.W.G., L.T.

## References

1. World_Health_Organisation. WHO World Malaria Report 2023. Geneva: World Health Organization; Licence: CC BY-NC-SA 30 IGO. 2023.

2. Rogerson SJ, Beeson JG, Laman M, Poespoprodjo JR, William T, Simpson JA, et al. Identifying and combating the impacts of COVID-19 on malaria. BMC Medicine. 2020;18(1):239.

3. van der Pluijm RW, Imwong M, Chau NH, Hoa NT, Thuy-Nhien NT, Thanh NV, et al. Determinants of dihydroartemisinin-piperaquine treatment failure in *Plasmodium falciparum* malaria in Cambodia, Thailand, and Vietnam: a prospective clinical, pharmacological, and genetic study. Lancet Infect Dis. 2019;19(9):952–61.

4. Imwong M, Dhorda M, Myo Tun K, Thu AM, Phyo AP, Proux S, et al. Molecular epidemiology of resistance to antimalarial drugs in the Greater Mekong subregion: an observational study. Lancet Infect Dis. 2020;20(12):1470–80.

5. Straimer J, Gandhi P, Renner KC, Schmitt EK. High prevalence of *Plasmodium falciparum* K13 mutations in Rwanda is associated with slow parasite clearance after treatment with artemether-lumefantrine. The Journal of infectious diseases. 2021;225(8):1411–4.

6. Balikagala B, Fukuda N, Ikeda M, Katuro OT, Tachibana S-I, Yamauchi M, et al. Evidence of artemisinin-resistant malaria in Africa. New England Journal of Medicine. 2021;385(13):1163–71.

7. Mihreteab S, Platon L, Berhane A, Stokes BH, Warsame M, Campagne P, et al. Increasing prevalence of artemisinin-resistant HRP2-negative malaria in Eritrea. N Engl J Med. 2023;389(13):1191–202.

8. Burrows JN, Duparc S, Gutteridge WE, Hooft van Huijsduijnen R, Kaszubska W, Macintyre F, et al. New developments in anti-malarial target candidate and product profiles. Malaria Journal. 2017;16(1):26.

9. Soucy TA, Smith PG, Milhollen MA, Berger AJ, Gavin JM, Adhikari S, et al. An inhibitor of NEDD8-activating enzyme as a new approach to treat cancer. Nature. 2009;458(7239):732-6.

10. Brownell JE, Sintchak MD, Gavin JM, Liao H, Bruzzese FJ, Bump NJ, et al. Substrate-assisted inhibition of ubiquitin-like protein-activating enzymes: the NEDD8 E1 inhibitor MLN4924 forms a NEDD8-AMP mimetic in situ. Mol Cell. 2010;37(1):102–11.

11. Hyer ML, Milhollen MA, Ciavarri J, Fleming P, Traore T, Sappal D, et al. A small-molecule inhibitor of the ubiquitin activating enzyme for cancer treatment. Nat Med. 2018;24(2):186–93.

12. Huang S-C, Adhikari S, Brownell JE, Calderwood EF, Chouitar J, D’Amore NR, et al. Discovery and optimization of pyrazolopyrimidine sulfamates as ATG7 inhibitors. Bioorganic & medicinal chemistry. 2020;28(19):115681.

13. Milhollen MA, Thomas MP, Narayanan U, Traore T, Riceberg J, Amidon BS, et al. Treatment-emergent mutations in NAEβ confer resistance to the NEDD8-activating enzyme inhibitor MLN4924. Cancer Cell. 2012;21(3):388–401.

14. Ferris J, Espona-Fiedler M, Hamilton C, Holohan C, Crawford N, McIntyre AJ, et al. Pevonedistat (MLN4924): mechanism of cell death induction and therapeutic potential in colorectal cancer. Cell Death Discovery. 2020;6(1):61.

15. Fu D-J, Wang T. Targeting NEDD8-activating enzyme for cancer therapy: developments, clinical trials, challenges and future research directions. Journal of Hematology & Oncology. 2023;16(1):87.

16. Xie SC, Metcalfe RD, Dunn E, Morton CJ, Huang S-C, Puhalovich T, et al. Reaction hijacking of tyrosine tRNA synthetase as a new whole-of-life-cycle antimalarial strategy. Science. 2022;376(6597):1074-9.

17. Xie SC, Wang Y, Morton CJ, Metcalfe RD, Dogovski C, Pasaje CFA, et al. Reaction hijacking inhibition of *Plasmodium falciparum* asparagine tRNA synthetase. Nat Commun. 2024;15(1):937.

18. Dogovski C, Xie SC, Burgio G, Bridgford J, Mok S, McCaw JM, et al. Targeting the cell stress response of *Plasmodium falciparum* to overcome artemisinin resistance. PLoS Biol. 2015;13(4):e1002132.

19. Fu Y, Tilley L, Kenny S, Klonis N. Dual labeling with a far red probe permits analysis of growth and oxidative stress in *P. falciparum*-infected erythrocytes. Cytometry A. 2010;77(3):253–63.

20. Bloch A, Coutsogeorgopoulos C. Inhibition of protein synthesis by 5’-sulfamoyladenosine. Biochemistry. 1971;10(24):4395–8.

21. Florini JR, Bird HH, Bell PH. Inhibition of protein synthesis in vitro and in vivo by nucleocidin, an antitrypanosomal antibiotic. J Biol Chem. 1966;241(5):1091–8.

22. Adhikari S, Calderwood EF, England DB, Gould AE, Harrison SJ, Huang S-C, et al. Atg7 inhibitors and the uses thereof. World Intellectual Property Organization. 2017;WO/2018/089786 PCT/US2017/061094.

23. Mandelbaum J, Rollins N, Shah P, Bowman D, Lee JY, Tayber O, et al. Identification of a lung cancer cell line deficient in atg7-dependent autophagy. Autophagy. 2015;(in press):00-.

24. Schulman BA, Harper JW. Ubiquitin-like protein activation by E1 enzymes: the apex for downstream signalling pathways. Nat Rev Mol Cell Biol. 2009;10(5):319–31.

25. Chen JJ, Tsu CA, Gavin JM, Milhollen MA, Bruzzese FJ, Mallender WD, et al. Mechanistic studies of substrate-assisted inhibition of ubiquitin-activating enzyme by adenosine sulfamate analogues. J Biol Chem. 2011;286(47):40867–77.

26. Baragaña B, Hallyburton I, Lee MC, Norcross NR, Grimaldi R, Otto TD, et al. A novel multiple-stage antimalarial agent that inhibits protein synthesis. Nature. 2015;522(7556):315-20.

27. Sanz LM, Crespo B, De-Cozar C, Ding XC, Llergo JL, Burrows JN, et al. *P. falciparum* in vitro killing rates allow to discriminate between different antimalarial mode-of-action. PloS one. 2012;7(2):e30949.

28. Supuran CT. Carbonic anhydrase inhibitors. Bioorg Med Chem Lett. 2010;20(12):3467–74.

29. Boddy A, Edwards P, Rowland M. Binding of sulfonamides to carbonic anhydrase: influence on distribution within blood and on pharmacokinetics. Pharm Res. 1989;6(3):203–9.

30. Angulo-Barturen I, Jimenez-Diaz MB, Mulet T, Rullas J, Herreros E, Ferrer S, et al. A murine model of falciparum-malaria by in vivo selection of competent strains in non-myelodepleted mice engrafted with human erythrocytes. PloS one. 2008;3(5):e2252.

31. Jimenez-Diaz MB, Mulet T, Gomez V, Viera S, Alvarez A, Garuti H, et al. Quantitative measurement of *Plasmodium-*infected erythrocytes in murine models of malaria by flow cytometry using bidimensional assessment of SYTO-16 fluorescence. Cytometry A. 2009;75(3):225–35.

32. Cowell AN, Istvan ES, Lukens AK, Gomez-Lorenzo MG, Vanaerschot M, Sakata-Kato T, et al. Mapping the malaria parasite druggable genome by using in vitro evolution and chemogenomics. Science. 2018;359(6372):191-9.

33. Luth MR, Gupta P, Ottilie S, Winzeler EA. Using in vitro evolution and whole genome analysis to discover next generation targets for antimalarial drug discovery. ACS Infect Dis. 2018;4(3):301–14.

34. Duffey M, Blasco B, Burrows JN, Wells TNC, Fidock DA, Leroy D. Assessing risks of *Plasmodium falciparum* resistance to select next-generation antimalarials. Trends in Parasitology. 2021;37(8):709–21.

35. Lv Z, Williams KM, Yuan L, Atkison JH, Olsen SK. Crystal structure of a human ubiquitin E1-ubiquitin complex reveals conserved functional elements essential for activity. J Biol Chem. 2018;293(47):18337–52.

36. Shen N, Guo L, Yang B, Jin Y, Ding J. Structure of human tryptophanyl-tRNA synthetase in complex with tRNATrp reveals the molecular basis of tRNA recognition and specificity. Nucleic Acids Res. 2006;34(11):3246–58.

37. Gamo FJ, Sanz LM, Vidal J, de Cozar C, Alvarez E, Lavandera JL, et al. Thousands of chemical starting points for antimalarial lead identification. Nature. 2010;465(7296):305-10.

38. Klonis N, Xie SC, McCaw JM, Crespo-Ortiz MP, Zaloumis SG, Simpson JA, et al. Altered temporal response of malaria parasites determines differential sensitivity to artemisinin. Proc Natl Acad Sci U S A. 2013;110(13):5157–62.

39. Straimer J, Gnadig NF, Witkowski B, Amaratunga C, Duru V, Ramadani AP, et al. K13-propeller mutations confer artemisinin resistance in *Plasmodium falciparum* clinical isolates. Science. 2014;Epub ahead of print.

40. Klonis N, Crespo-Ortiz MP, Bottova I, Abu-Bakar N, Kenny S, Rosenthal PJ, et al. Artemisinin activity against *Plasmodium falciparum* requires hemoglobin uptake and digestion. Proc Natl Acad Sci U S A. 2011;108(28):11405–10.

41. Johnson JD, Dennull RA, Gerena L, Lopez-Sanchez M, Roncal NE, Waters NC. Assessment and continued validation of the malaria SYBR green I-based fluorescence assay for use in malaria drug screening. Antimicrob Agents Chemother. 2007;51(6):1926–33.

42. Aguiar AC, Pereira DB, Amaral NS, De Marco L, Krettli AU. *Plasmodium vivax* and *Plasmodium falciparum ex vivo* susceptibility to anti-malarials and gene characterization in Rondonia, West Amazon, Brazil. Malar J. 2014;13:73.

43. Reader J, Botha M, Theron A, Lauterbach SB, Rossouw C, Engelbrecht D, et al. Nowhere to hide: interrogating different metabolic parameters of Plasmodium falciparum gametocytes in a transmission blocking drug discovery pipeline towards malaria elimination. Malar J. 2015;14:213.

44. Reader J, van der Watt ME, Birkholtz LM. Streamlined and robust stage-specific profiling of gametocytocidal compounds against *Plasmodium falciparum*. Front Cell Infect Microbiol. 2022;12:926460.

45. Delves MJ, Straschil U, Ruecker A, Miguel-Blanco C, Marques S, Dufour AC, et al. Routine in vitro culture of *P. falciparum* gametocytes to evaluate novel transmission-blocking interventions. Nat Protoc. 2016;11(9):1668–80.

46. Miglianico M, Bolscher JM, Vos MW, Koolen KJM, de Bruijni M, Rajagopal DS, et al. Assessment of the drugability of initial malaria infection through miniaturized sporozoite assays and high-throughput screening. Commun Biol. 2023;6(1):216.

47. Yang ASP, van Waardenburg YM, van de Vegte-Bolmer M, van Gemert GA, Graumans W, de Wilt JHW, et al. Zonal human hepatocytes are differentially permissive to *Plasmodium falciparum* malaria parasites. Embo J. 2021;40(6):e106583.

48. Yang JJ, Milton MN, Yu S, Liao M, Liu N, Wu JT, et al. P-glycoprotein and breast cancer resistance protein affect disposition of tandutinib, a tyrosine kinase inhibitor. Drug Metab Lett. 2010;4(4):201–12.

49. Jimenez-Diaz MB, Mulet T, Viera S, Gomez V, Garuti H, Ibanez J, et al. Improved murine model of malaria using Plasmodium falciparum competent strains and non-myelodepleted NOD-scid IL2Rgammanull mice engrafted with human erythrocytes. Antimicrob Agents Chemother. 2009;53(10):4533–6.

50. Aragão D, Aishima J, Cherukuvada H, Clarken R, Clift M, Cowieson NP, et al. MX2: a high-flux undulator microfocus beamline serving both the chemical and macromolecular crystallography communities at the Australian Synchrotron. J Synchrotron Radiat. 2018;25(Pt 3):885–91.

51. Evans PR, Murshudov GN. How good are my data and what is the resolution? Acta Crystallogr D. 2013;69:1204–14.

52. Winn MD, Ballard CC, Cowtan KD, Dodson EJ, Emsley P, Evans PR, et al. Overview of the CCP4 suite and current developments. Acta Crystallogr D. 2011;67:235–42.

53. Mccoy AJ, Grosse-Kunstleve RW, Adams PD, Winn MD, Storoni LC, Read RJ. Phaser crystallographic software. Journal of Applied Crystallography. 2007;40:658–74.

54. Adams PD, Afonine PV, Bunkoczi G, Chen VB, Davis IW, Echols N, et al. PHENIX: a comprehensive Python-based system for macromolecular structure solution. Acta Crystallogr D. 2010;66:213–21.

55. Emsley P, Lohkamp B, Scott WG, Cowtan K. Features and development of Coot. Acta Crystallographica Section D-Biological Crystallography. 2010;66:486–501.

56. Moriarty NW, Grosse-Kunstleve RW, Adams PD. Electronic ligand builder and optimization workbench (eLBOW): A tool for ligand coordinate and restraint generation. Acta Crystallographica Section D: Biological Crystallography. 2009;65:1074–80.

57. Mujumdar P, Bua S, Supuran CT, Peat TS, Poulsen SA. Synthesis, structure and bioactivity of primary sulfamate-containing natural products. Bioorg Med Chem Lett. 2018;28(17):3009–13.

58. Hall J. A simple model for determining affinity from irreversible thermal shifts. Protein Sci. 2019;28(10):1880–7.

