## Supplementary Information for "A potent and selective reaction hijacking inhibitor of *Plasmodium falciparum* tyrosine tRNA synthetase exhibits single dose oral efficacy *in vivo*"

**The PDF file includes:**

Supplementary Figures 1 to 8
Supplementary Tables 1 to 12
Chemistry Materials and Methods
Supplementary References

#### Supplementary Figures

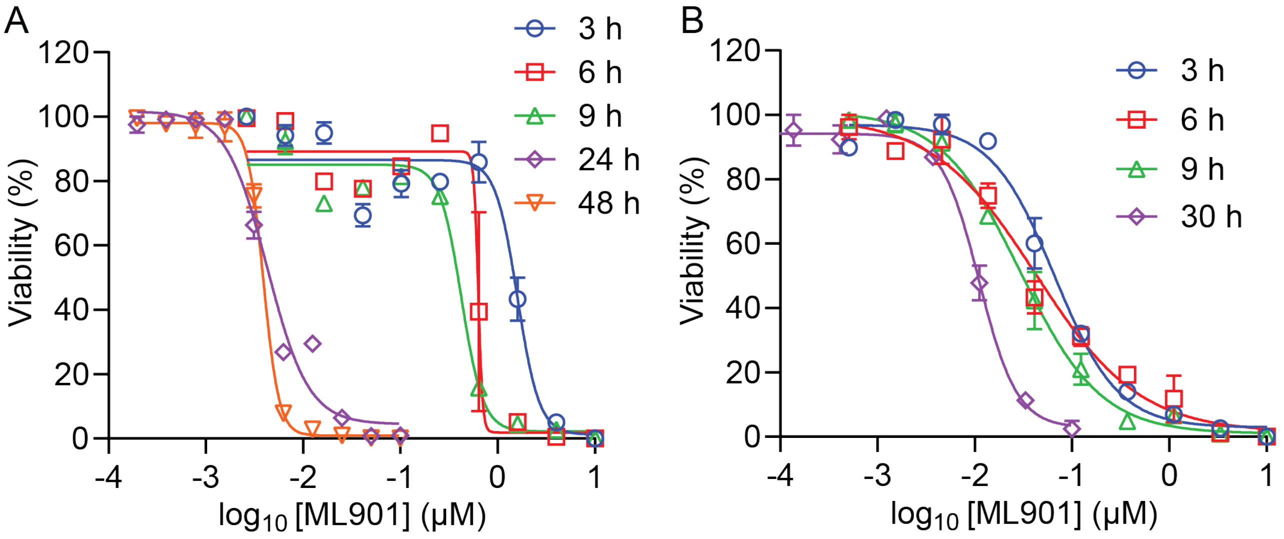

**Supplementary Figure 1. Exposure-time dependent responses of trophozoite and schizont stage parasite to short pulses of ML901**

(A,B) A tightly synchronized culture of CAM 3.II Rev parasites (>70% of parasites within a 5-h time window) was subjected to pulses of ML901 for 3 h, 6 h, 9 h, 24 h, or continued exposure for 48 h (A) or 30 h (B), initiated at (A) trophozoite (25-30 h.p.i.) and (B) schizont (43-48 h.p.i.) stages. Flow cytometric analysis of Syto-61-labelled parasites in the cycle after the initiation of treatment assesses cell viability.

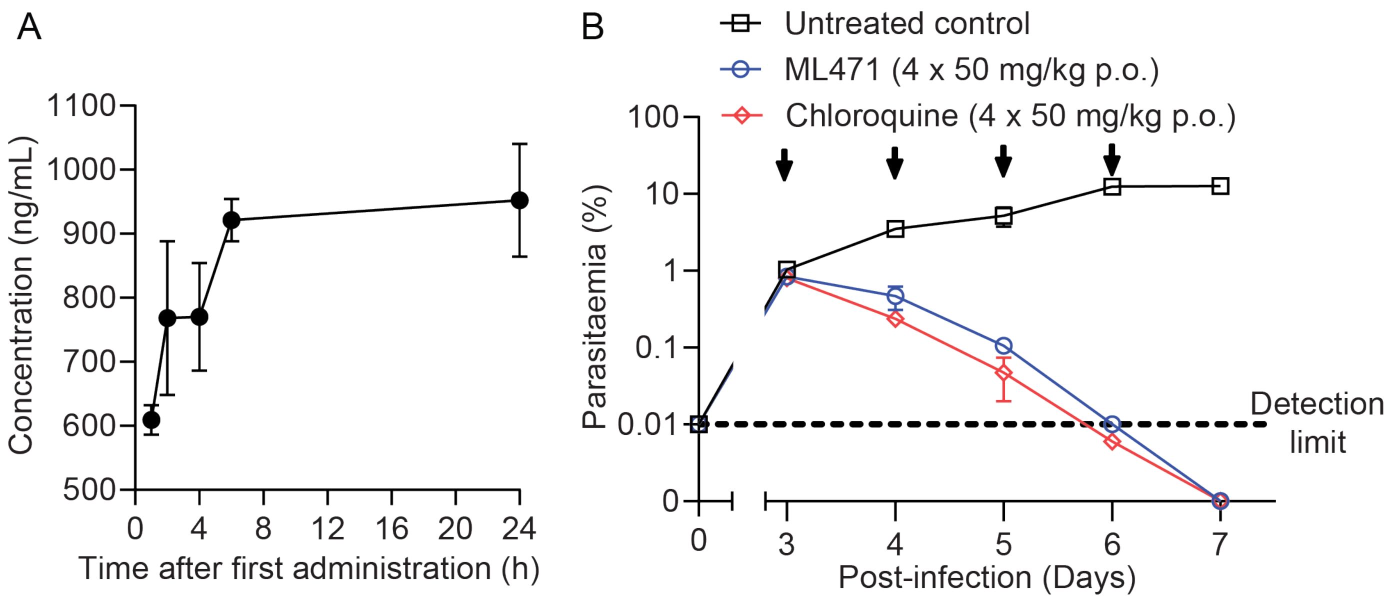

**Supplementary Figure 2. Pharmacokinetics profile and efficacy of ML471 in treating *P. falciparum* infected mice.**

(A) Pharmacokinetics profile (in blood), for SCID mice engrafted with human RBCs infected with *P. falciparum*, over the first day following treatment with ML471 at 50 mg/kg p.o. See Suppl Table S8 for pharmacokinetics values. (B) Therapeutic efficacy of ML471 in the SCID mouse *P. falciparum* model, dosed with ML471 for 4 days at 50 mg/kg p.o. per day (arrows), initiated on Day 3 post-infection. The chloroquine data are from [1].

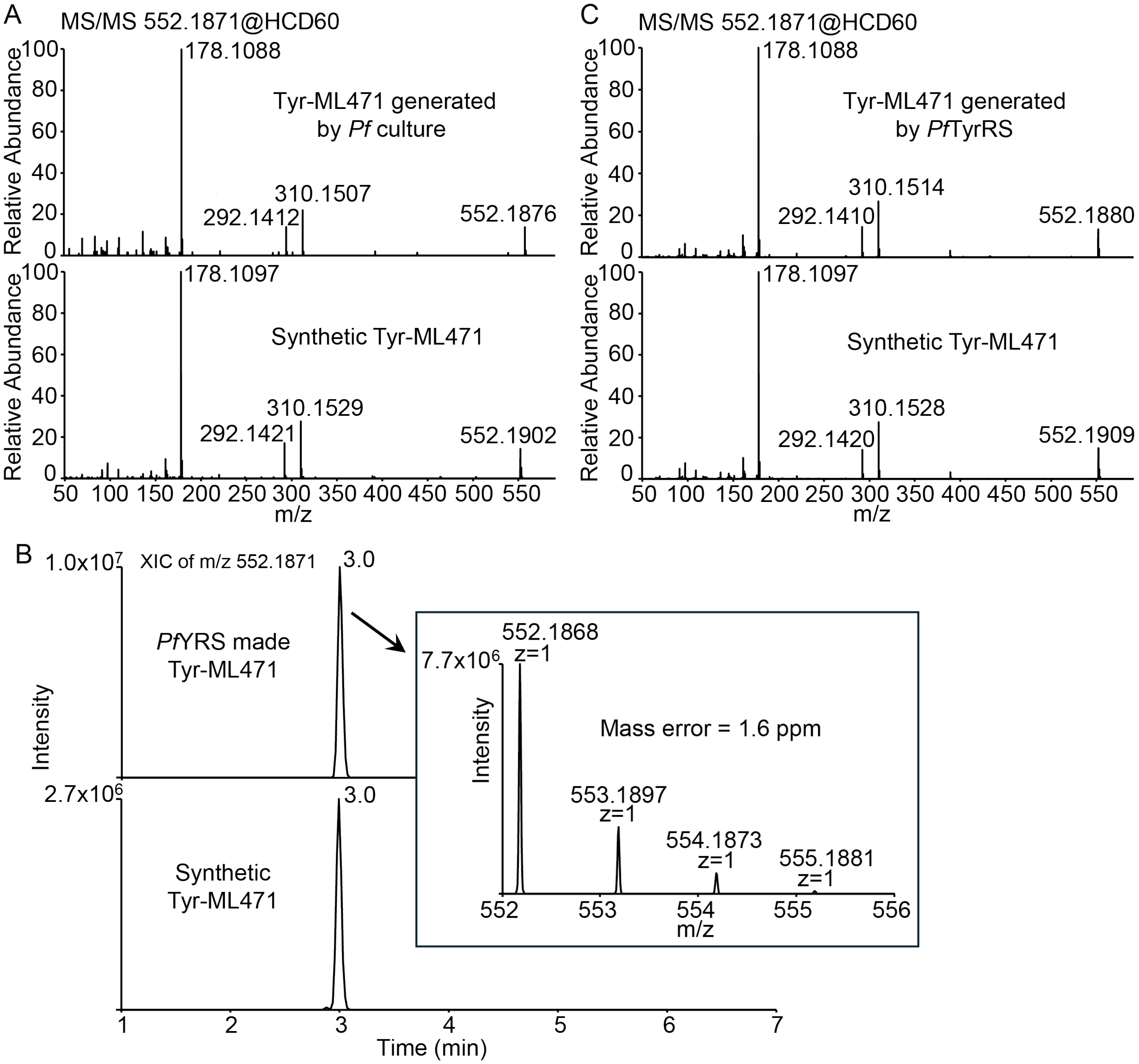

**Supplementary Figure 3. Identification of Tyr-ML471 adduct made by *P. falciparum* culture and *Pf*TyrRS enzyme.**

(A) MS/MS analysis of the Tyr-ML471 adduct made by *P. falciparum* following treatment with ML471 (1 μM) for 2 h (upper panel); and the synthetic conjugate at 0.2 μM (lower panel). (B,C) *Pf*TyrRS (1 μM) was incubated with ML471 (10 μM), ATP (10 μM), tyrosine (20 μM) and 4 μM *Pf*tRNA$^{Tyr}$ for 1 h at 37°C. Following protein denaturation and precipitation, the supernatant was subjected to LCMS analysis. (B) The extracted ion chromatograms of Tyr-ML471 adduct made by *Pf*TyrRS (upper panel); and the synthetic conjugate at 1 μM (lower panel). The inset shows the MS analysis of the enzyme-generated Tyr-ML471. (C) MS/MS analysis of the enzyme-generated Tyr-ML471 (upper panel) and the synthetic conjugate at 1 μM (lower panel).

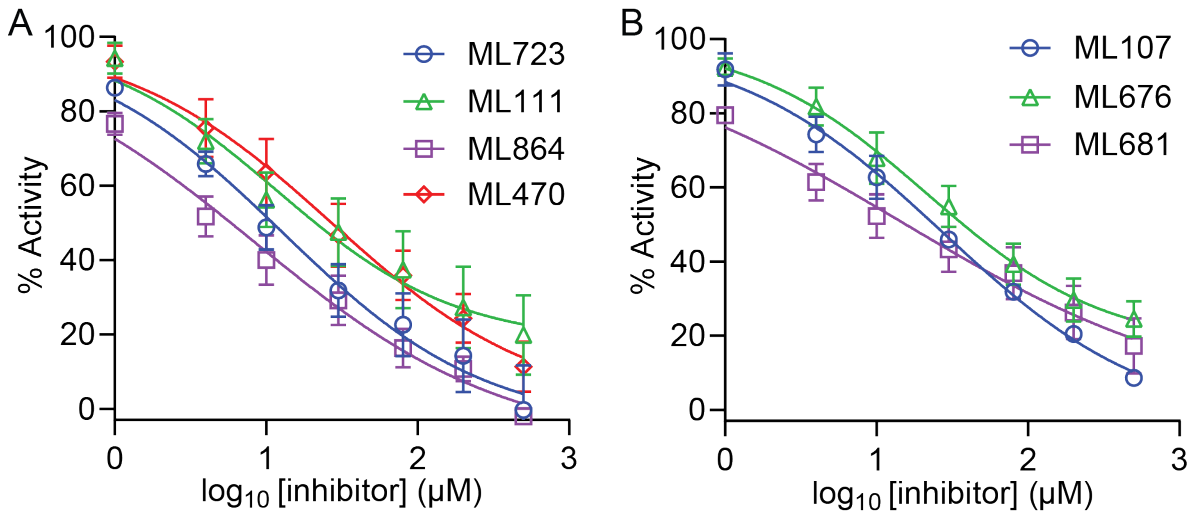

**Supplementary Figure 4. Kinase GLO assays for ML901 and derivatives.**
Effects of increasing concentrations of ML723, ML111, ML864, ML470 (A) and ML107, ML676, ML681 (B) on ATP consumption by *Pf*TyrRS. The reaction conditions are: *Pf*TyrRS (25 nM), ATP (10 µM), tyrosine (200 µM), cognate $tRNA^{Tyr}$ (4.8 µM) and pyrophosphatase (1 unit/mL). Incubations were at 37°C for 1 h. Data are mean ± SEM from three independent experiments

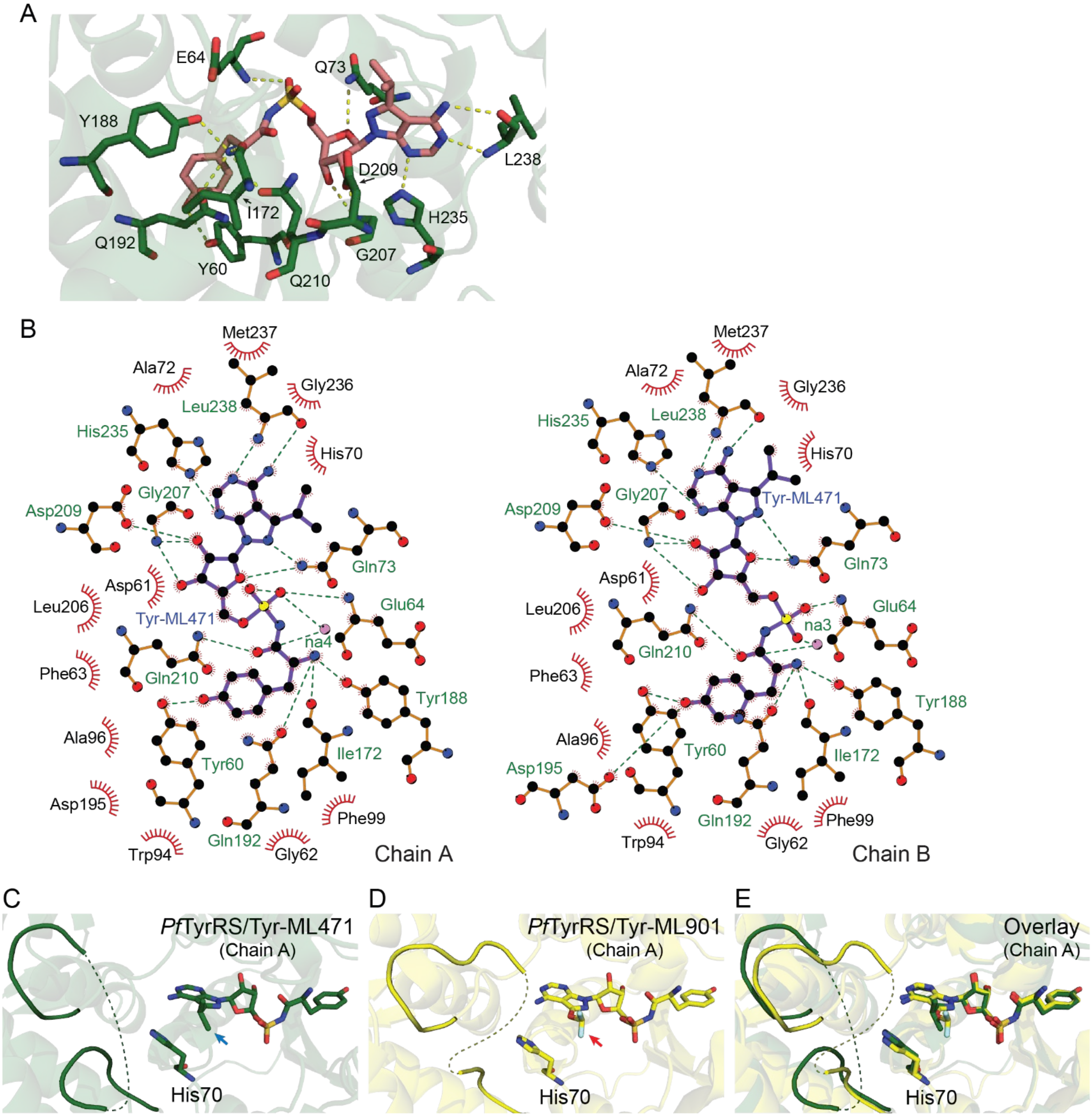

**Supplementary Figure 5. Comparison of the active site architecture of *Pf*TyrRS in complex with Tyr-ML471 or Tyr-ML901 (7ROS).** (A) Inhibitor/active site interactions for the A-chain of *Pf*TyrRS with bound Tyr-ML471. (B) LigPlots of interacting residues for the A- and B-chains of *Pf*TyrRS with bound Tyr-ML471. (C) A-chain of Tyr-ML471-bound *Pf*TyrRS showing the poses adopted by the ML471 isopropyl (aqua arrow) and His70, which are incompatible with a structured KMSKS loop. (D) A-chain of Tyr-ML901-bound *Pf*TyrRS (7ROS) illustrating the ML901 difluoromethoxy group (red arrow) and the His70 conformation. The KMSKS loop is not resolved. (E) Overlay of the A-chains of Tyr-ML471- and Tyr-ML901-bound *Pf*TyrRS.

#### Supplementary Tables

**Table S1. The median lethal dose ($LD_{50}$) values of different asexual blood stages after exposure of ML901.**

| Drug pulse time (h) | $LD_{50}$ (nM) | |
|---|---|---|
| | Troph | Schizont |
| 3 | 1700 ± 460 | 88 ± 1 |
| 6 | 680 ± 48 | 36 ± 6 |
| 9 | 290 ± 150 | 21 ± 12 |
| 24 | 4.0 ± 0.0 | N/D |
| 48 (trophozoite), 30 (schizont), continuous | 3.9 ± 0.8 | 8.5 ± 0.7 |

**Table S2**. ***P. vivax* and *P. falciparum* ex vivo drug susceptibility.**

| Compound | Brazilian field isolates | |
|---|---|---|
| | *P. falciparum* | *P. vivax* |
| | Median $EC_{50}$ (nM) (Range) [Number of isolates] | |
| ML471 | 4.0 (2.9-4.2) [n = 7] | 6.7 (4.1-59) [n = 10] |
| Artesunate | 1.8 (0.9-2.1) [n = 7] | 1.5 (0.79-3.1) [n = 10] |
| Chloroquine | 276 (152-903) [n = 7] | 133 (105-204) [n = 10] |

**Table S3**. **Activity against immature (>90% stage II/III) and mature (>95% stage V) stage gametocytes (Pf 3D7-pfs16-CBG99) for ML901, ML471 and antimalarial controls.**

| Compound | *Early stage gametocytes* $IC_{50}$ (nM) | *Mature stage gametocytes* $IC_{50}$ (nM) |
|---|---|---|
| ML901 | 310 ± 60 (n = 3) | 1660 ± 140 (n = 3) |
| ML471 | 112 ± 8 (n = 3) | 392 ± 14 (n = 3) |
| MMV390048 | 138 ± 8 (n=3) | 150 ± 20 (n=3) |
| MB Control | 62 ± 20 (n=3) | 353 ± 63 (n=3) |

**Table S4**. **Activity against *P. falciparum* schizonts in primary hepatocytes.**

| Compound | *P. falciparum liver stage schizont (NF175)* $IC_{50}$ (nM) | *Primary hepatocytes* $IC_{50}$ (nM) | *P. falciparum liver stage schizont (NF135)* $IC_{50}$ (nM) | *Primary hepatocytes* $IC_{50}$ (nM) |
|---|---|---|---|---|
| ML901 | 13 ± 5 nM (n = 3)* | >10,000 (n = 3) | 2.9 ± 0.9 (n = 3) | > 100 (n = 3) |
| ML471 | 2.8 ± 1.5 (n = 3) | >10,000 (n = 3) | 5.5 ± 2.2 (n = 3) | > 100 (n = 3) |
| Atovaquone | 7.0 ± 3.9 (n = 3) | >1,000 (n = 3) | N.A. | N.A. |
| MMV390048 | N.A. | N.A. | 23.3 ± 13.3 (n = 3) | >100 (n = 3) |
| * Data from [1] | | | | |

**Table S5**. **Activity against transmissible gametes.**

| Compound | Dual gamete formation assay (male) $IC_{50(48h)}$ (nM)* | Dual gamete formation assay (female) $IC_{50(48h)}$ (nM)* |
|---|---|---|
| ML901 | 130 ± 10 (n = 9)* | 4,700 ± 300 (n = 9)* |

| | | |
|---|---|---|
| **ML471** | 49 ± 3 (n = 4) | 260 ± 10 (n = 4) |
| **Cabamiquine (DDD498)** | 2.3 ± 0.4 (n = 5) | 2.2 ± 0.1 (n = 5) |
| * Data from [1]. Data for the DDD498 control is very similar to a previous report [2], validating the assay. | | |

**Table S6. Parasite Reduction Ratio**

| **Compound** | **Dose** | **Lag phase (h)** | **Slope** | **R** | **Log PRR** | **PCT99.9% (h)** |
|---|---|---|---|---|---|---|
| **ML471** | 10 x $IC_{50}$ | 0 | -0.09 | 0.96 | 4.1 | 32 |
| **Controls** | | | | | | |
| **Pyrimethamine** | 10 x $IC_{50}$ | 24 | -0.078 | 0.98 | 3.7 | 56 |
| **Chloroquine*** | 10 x $IC_{50}$ | 0 | -0.094 | 0.99 | 4.5 | 33.8 |
| **Atovaquone*** | 10 x $IC_{50}$ | 48 | -0.060 | 0.99 | 2.9 | 90 |
| **Artemisinin*** | 10 x $IC_{50}$ | 0 | nd | nd | >4.8 | <24 |
| *Data from [3]. | | | | | | |

**Table S7. Pharmacological and ADME characterization of ML471.**
For PK studies, rats (n = 2) were dosed with ML471 at 1 mg/kg i.v. or 10, 20 or 25 mg/kg p.o. and plasma and blood samples were collected for analysis. For blood to plasma ratio, rats (n = 2) were dosed with ML471 at 1 mg/kg and plasma and blood samples were collected for analysis.

| **PK analysis** | | | | | | | | | | |
|---|---|---|---|---|---|---|---|---|---|---|
| **Route** | **Dose** | **Matrix** | **AUC (INF)** | $T_{max}$ | $C_{max}$ | $T_{1/2z}$ | **CL** | **Vss** | **Vz** | **%F** |
| | **mg/kg** | | **(h*mM)** | **(h)** | **(nM)** | **(h)** | **(L/h/kg)** | **(L/kg)** | **(L/kg)** | |
| IV | 1 | Blood | 21.00 | 0.17 | 1180 | 36.6 | 0.138 | 6.25 | 6.54 | |
| IV | 1 | Plasma | 0.70 | 0.08 | 1540 | 2.22 | 3.84 | 2.79 | 12.3 | |
| PO | 1 | Blood | 1.95 | 16.0 | 39.3 | 23.2 | | | | 9.29 |
| PO | 1 | Plasma | 0.194 | 0.5 | 19.1 | 8.78 | | | | 27.7 |
| PO | 10 | Blood | 13.15 | 6.0 | 261 | 29.5 | | | | 9.56 |
| PO | 10 | Plasma | 1.74 | 0.38 | 196 | 9.15 | | | | 25.2 |
| PO | 25 | Blood | 29.88 | 8.0 | 669 | 30.5 | | | | 8.72 |
| PO | 25 | Plasma | 2.84 | 0.75 | 393 | 3.57 | | | | 16.4 |

| **Blood to plasma ratio** | | | | | | | |
|---|---|---|---|---|---|---|---|
| Time (h) | 0.083 | 0.25 | 0.5 | 1 | 2 | 4 | 8 |
| Mean blood to plasma ratio | 0.8 | 1.5 | 2.1 | 5.5 | 17.8 | 48.9 | 113.9 |

| **ADME data** | | | |
|---|---|---|---|
| Compound | AlogP/ TSPA | Solubility (μM) | Caco2 $P_{app}$ ($x10^{-6}$ cm/s) (A to B; B to A) |
| ML471 | -1.18/ 189 | >100 | 7.6, 0.17, 1.3. |

**Table S8. Pharmacological parameters for ML471 in the SCID mouse model.**
Mice were dosed orally with four daily doses of 50 mg/kg. Samples were collected from day 3, at 1, 2, 4, 6 and 24 h after the first administration. In a separate experiment, mice were dosed orally with one dose of 100 or 200 mg/kg on day 3 after infection. Samples were collected from day 3, at 1, 2, 4, 6, 24, 48, 72, 96 and 120 h after the first administration.

| **SCID mouse study 4 days x 50 mg/kg p.o.** | | | | | | |
|---|---|---|---|---|---|---|
| **Mouse** | Dose | **$T_{max}$ [h]** | **$C_{max}$ [ng/mL]** | **$C_{max}$ /dose [(ng/mL)/(mg/kg)]** | **AUC0-24h [h*ng/mL]** | **AUC0-24h/dose [(h*ng/mL)/(mg/kg)]** |
| **M1** | 50 | 6 | 888 | 17.8 | 19,586 | 392 |
| **M2** | 50 | 24 | 1,040 | 20.8 | 22,572 | 451 |
| **N** | | 2 | 2 | 2 | 2 | 2 |
| **Mean** | | 15 | 964 | 19.3 | 21,100 | 422 |
| **SD** | | 12.7 | 107 | 2.2 | 2,100 | 42 |
| **SCID mouse study 1 x 100 or 200 mg/kg p.o.** | | | | | | |
| **Mouse** | Dose | **$T_{max}$ [h]** | **$C_{max}$ [ng/mL]** | **$C_{max}$ /dose [(ng/mL)/(mg/kg)]** | **AUC0-120h [h*ng/mL]** | **AUC0-120h/dose [(h*ng/mL)/(mg/kg)]** |
| **M1** | 100 | 4 | 3,100 | 31 | 239,000 | 2,390 |
| **M2** | 100 | 6 | 3,600 | 36 | 260,000 | 2,600 |
| **N** | | 2 | 2 | 2 | 2 | 2 |
| **Mean** | | 5 | 3,350 | 33.5 | 250,000 | 2,500 |
| **SD** | | 1.4 | 354 | 3.54 | 14,500 | 145 |
| **M3** | 200 | 4 | 3,340 | 16.7 | 171,000 | 855 |
| **M4** | 200 | 4 | 4,180 | 20.9 | 259,000 | 1,290 |
| **N** | | 2 | 2 | 2 | 2 | 2 |
| **Mean** | | 4 | 3,760 | 18.8 | 215,000 | 1,070 |
| **SD** | | 0 | 594 | 2.97 | 62,000 | 310 |

**Table S9. Summary of resistance selection.**
For ML901, Dd2-B2 parasites were subjected to pressure at 3 x $IC_{50}$. Mean $IC_{50}$ = 2.6 ± 0.05 nM. For ML471, Dd2-B2 parasites were subjected to a single-step selection at 10 x $IC_{50}$. Mean $IC_{50}$ and $IC_{90}$ values were 1.45 and 1.99 nM.

| **ML901** | | |
|---|---|---|
| Number of parasites in the selection | 10^7 | 10^8 |
| Day of recrudescence | 12 (1/3), 14 (2/3) | 14 (3/3) |
| $IC_{50}$ fold shift (clones) | 2-3 x $IC_{50}$ | 3-6 x $IC_{50}$ |
| Minimum Inoculum for Resistance (MIR) | ≤ 10^7 | |

| **ML471** | |
|---|---|
| Number of parasites in the selection | 2x10^5 |
| Day of recrudescence | 18 (29/96 wells) |
| $IC_{50}$ shift | 9-16 x$IC_{50}$ |
| Minimum Inoculum for Resistance (MIR) | 7.1 x10^5 |

**Table S10. Copy Number Variants (CNV) in ML901-selected sample**

CNV amplification was found in sample F1 (BULK_JOS_1E7_F1), F3 (BULK_JOS_1E8_F3), F1-F6 (F1_F6_KS, a clone from the BULK 1E8 F1 selection) and FL1 (FL1_1E8) at different factors. F1 exhibits a lower increase at ~1.8 across a wider range, whereas F3, F1-F6 and FL1 show a ~2x increase only for the second half of the segment. Strikingly, three separate genome amplification boundaries were observed, with each including the tyrosine tRNA ligase (PF3D7_0807900).

| Gene ID | Gene Name | Fold gain in copy number on Chromosome 8 | | | |
|---|---|---|---|---|---|
| | | F1 | F3 | F1_F6 | FL1 |
| PF3D7_0806300 | ferlin-like protein, putative | 1.80 | 1.00 | 1.00 | 1.00 |
| PF3D7_0806400 | UDP-N-acetylglucosamine transferase subunit ALG13, putative | 1.80 | 1.00 | 1.00 | 1.00 |
| PF3D7_0806500 | DnaJ protein, putative | 1.80 | 1.00 | 1.00 | 1.00 |
| PF3D7_0806600 | kinesin-like protein, putative | 1.80 | 1.00 | 1.00 | 1.00 |
| PF3D7_0806700 | conserved Plasmodium membrane protein, unknown function | 1.80 | 1.00 | 1.00 | 1.00 |
| PF3D7_0806800 | V-type proton ATPase subunit a, putative | 1.80 | 1.00 | 1.00 | 1.00 |
| PF3D7_0806900 | conserved protein, unknown function | 1.80 | 1.00 | 1.00 | 1.00 |
| PF3D7_0807000 | YEATS domain-containing protein, putative | 1.80 | 1.00 | 1.00 | 1.79 |
| PF3D7_0807100 | DNA helicase PSH3 | 1.80 | 1.00 | 1.00 | 1.79 |
| PF3D7_0807200 | conserved Plasmodium membrane protein, unknown function | 1.67 | 2.24 | 2.54 | 2.32 |
| PF3D7_0807300 | ras-related protein Rab-18 | 1.67 | 2.24 | 2.54 | 2.32 |
| PF3D7_0807400 | coenzyme Q-binding protein COQ10 homolog, mitochondrial | 1.67 | 2.24 | 2.54 | 2.32 |
| PF3D7_0807500 | proteasome subunit alpha type-6, putative | 1.67 | 2.24 | 2.54 | 2.32 |
| PF3D7_0807600 | conserved Plasmodium protein, unknown function | 1.67 | 2.24 | 2.54 | 2.32 |
| PF3D7_0807700 | serine protease DegP | 1.67 | 2.24 | 2.54 | 2.32 |
| PF3D7_0807800 | 26S proteasome regulatory subunit RPN10, putative | 1.67 | 2.24 | 2.54 | 2.32 |
| **PF3D7_0807900** | **tyrosine tRNA ligase** | **1.67** | **2.24** | **2.54** | **2.32** |
| PF3D7_0808000 | conserved Plasmodium protein, unknown function | 1.00 | 2.24 | 2.54 | 2.32 |
| PF3D7_0808100 | AP-3 complex subunit delta, putative | 1.00 | 2.24 | 2.54 | 2.32 |
| PF3D7_0808200 | plasmepsin X | 1.00 | 2.24 | 2.54 | 2.32 |
| PF3D7_0808300 | ubiquitin regulatory protein, putative | 1.00 | 2.24 | 2.54 | 2.32 |
| PF3D7_0808400 | coatomer subunit epsilon, putative | 1.00 | 2.24 | 2.54 | 2.32 |

**Table S11. Copy Number Variants (CNV) in ML471-selected sample.**

CNV amplification within a 57 kb segment on chromosome 8, by ~3.8 and 4-fold in samples C2 and E2, respectively. These samples correspond to wells seeded at 2x10^5 parasites that recrudesced following selection at 10x $IC_{50}$. Note that both wells had different 5' amplification breakpoints and thus were independent events that shared the same region of amplification at the 3' end. Both wells shared a ~4-fold amplification of tyrosine tRNA ligase, consistent with a slightly higher $IC_{50}$ increase compared with ML901-selected ~2-fold amplifications described above.

| Gene ID | Gene Name | Fold gain in copy number on Chromosome 8 | |
|---|---|---|---|
| | | C2 | E2 |

| | | | |
|---|---|---|---|
| PF3D7_0807000 | YEATS domain-containing protein, putative | 3.48 | 1.000 |
| PF3D7_0807100 | DNA helicase PSH3 | 3.48 | 1.000 |
| PF3D7_0807200 | conserved Plasmodium membrane protein, unknown function | 3.80 | 1.000 |
| PF3D7_0807300 | ras-related protein Rab-18 | 3.80 | 1.00 |
| PF3D7_0807400 | coenzyme Q-binding protein COQ10 homolog, mitochondrial | 3.80 | 1.00 |
| PF3D7_0807500 | proteasome subunit alpha type-6, putative | 3.80 | 1.00 |
| PF3D7_0807600 | conserved Plasmodium protein, unknown function | 3.80 | 1.00 |
| PF3D7_0807700 | serine protease DegP | 3.80 | 4.04 |
| PF3D7_0807800 | 26S proteasome regulatory subunit RPN10, putative | 3.80 | 4.04 |
| **PF3D7_0807900** | **tyrosine tRNA ligase** | **3.80** | **4.04** |
| PF3D7_0808000 | conserved Plasmodium protein, unknown function | 3.80 | 4.04 |
| PF3D7_0808100 | AP-3 complex subunit delta, putative | 3.80 | 4.04 |
| PF3D7_0808200 | plasmepsin X | 3.80 | 4.04 |
| PF3D7_0808300 | ubiquitin regulatory protein, putative | 3.80 | 4.04 |
| PF3D7_0808400 | coatomer subunit epsilon, putative | 3.80 | 4.04 |
| PF3D7_0808500 | Plasmodium RNA of unknown function RUF6 | 3.80 | 4.04 |

**Table S12. X-ray diffraction data collection and refinement statistics.**

| | ***Pf*YRS/Tyr-ML471** |
|---|---|
| **Data collection** | |
| Space group | C 2 |
| Wavelength (Å) | 0.9537 |
| Number of images | 3600 |
| Oscillation range per image (˚) | 0.1 |
| Detector | Eiger 16M |
| *Cell dimensions* | |
| a, b, c (Å) | 138.21, 47.27, 140.512 |
| α, β, γ (˚) | 90, 94.63, 90 |
| Resolution (Å) | 46.68 - 1.80 (1.82 - 1.80) |
| Rsym | 0.086 (0.765) |
| Rmeas | 0.102 (0.899) |
| Rpim | 0.054 (0.470) |
| CC1/2 | 0.998 (0.838) |
| I/σ(I) | 9.4 (1.2) |
| Total observations | 567540 (31557) |
| Unique reflections | 84432 (4481) |

| | |
|---|---|
| Completeness (%) | 99.9 (100.0) |
| Multiplicity | 6.7 (7.0) |
| Wilson B factor (Å) | 32.5 |
| **Refinement** | |
| Resolution (Å) | 46.68 - 1.80 (1.82 - 1.80) |
| Reflections used in refinement | 84394 (2835) |
| Rfree reflections | 4177 (136) |
| $R_{work}$ | 0.1830 (0.3172) |
| $R_{free}$ | 0.2198 (0.3517) |
| Protein molecules in asymmetric unit | 2 |
| Total nonhydrogen atoms | 6159 |
| Protein | 5683 |
| Ligand/ion | 96 |
| Solvent | 380 |
| Mean B factor ($Å^2$) | 50.31 |
| Protein | 50.50 |
| Ligand/ion | 39.63 |
| Solvent | 50.14 |
| RMS deviations | |
| Bond lengths (Å) (outliers > 4 $\sigma$ ) | 0.017 |
| Bond angles (°) (outliers > 4 $\sigma$ ) | 1.52 |
| Rotamer outliers | 1.09 |
| Clashscore | 6.33 |
| Cβ outliers | 0 |
| Molprobity score | 1.35 |
| *Ramachandran Plot* | |
| Favoured (%) | 98.55 |
| Allowed (%) | 1.45 |
| Outliers (%) | 0 |

**Supplementary Chemistry Materials and Methods**

*Chemical abbreviations*

Boc: *tert*-Butoxycarbonyl
DIPEA: *N,N*-Diisopropylethylamine
DMA: *N,N*-Dimethylacetamide
DMAP: 4-Dimethylaminopyridine
DMF: *N,N*-Dimethylformamide
DMSO: Dimethyl sulfoxide
EDCI: 1-Ethyl-3-(3-dimethylaminopropyl) carbodiimide
HPLC: High-performance liquid chromatography
LC-MS: Liquid chromatography-mass spectrometry
NIS: *N*-Iodosuccinimide
NMR: Nuclear magnetic resonance spectroscopy
*p*-TsOH: *p*-Toluenesulfonic acid
TFA: Trifluoroacetic acid
THF: Tetrahydrofuran
TPPTS: Triphenylphosphine-3,3′,3″-trisulfonic acid trisodium salt
Tyr: Tyrosine

*General information*

Reagents were purchased from Sigma-Aldrich, Merck, Fisher Scientific, and Combi-Blocks, and were used without further purification. Anhydrous conditions: glassware was dried at >130 °C for >12 h, assembled hot, and purged with nitrogen ($N_2$) gas where suitable. Reduced pressure/*in vacuo* implies 900 to 50 mbar under rotary evaporation at 40–50 °C.

Liquid chromatography-mass spectrometry (LC-MS) data were documented on an Agilent™ 1260 Infinity II LC System with a Diode Array HS (G7117C) UV-Visible detector coupled to a triple quadrupole mass detector. (analytical column: Pursuit XR C18 100 Å, 3 µm, 2 mm × 50 mm; solvent gradient: 5–95% acetonitrile in 0.05% v/v aqueous trifluoroacetic acid; flow rate: 0.4 ml min$^{-1}$; elution run time: 14 minutes). The charge of the ion specifies that the detection is positive; for instance, $[M+H]^+$ denotes positive-ion detection.

Analytical thin-layer chromatography (TLC) was carried out on Merck Silica Gel 60 $F_{254}$-precoated aluminum plates (0.2 mm) and observed using UV irradiation (254 nm and 280 nm). High-temperature reactions were carried out in DrySyn heating blocks.

Proton ($^1$H) NMR spectra were recorded on a BrukerDRX400 spectrometer operating at 400 MHz for proton nuclei. Deuterated solvents ($CDCl_3$, $CD_3OD$ and DMSO-$d_6$) were obtained from Sigma-Aldrich. $^1$H chemical shifts are stated in parts per million (ppm). Spectroscopic chemical shifts were calibrated to residual solvent peaks ($^1$H: $CHCl_3$ 7.26 ppm, methanol 3.31ppm, dimethyl sulfoxide 2.50 ppm). The multiplicities are labelled as either a singlet (s), doublet (d), triplet (t), doublet of doublets (dd), or multiplet (m).

For preparative HPLC purification, samples were injected onto a Phenomenex Luna® C8 column (5 µM particle size, 100 Å, 150 x 21.2 mm) and analysed on Agilent Technologies 1260 Infinity II HPLC system equipped with a photodiode array detector, and a preparative fraction collector (G1364E), (Solvent gradient: 2–35% acetonitrile in 0.05% v/v aqueous trifluoroacetic acid; flow rate: 15 ml min$^{-1}$; elution run time: 40 minutes).

#### Synthesis and Characterisation of *ML471 and Tyr-ML471*

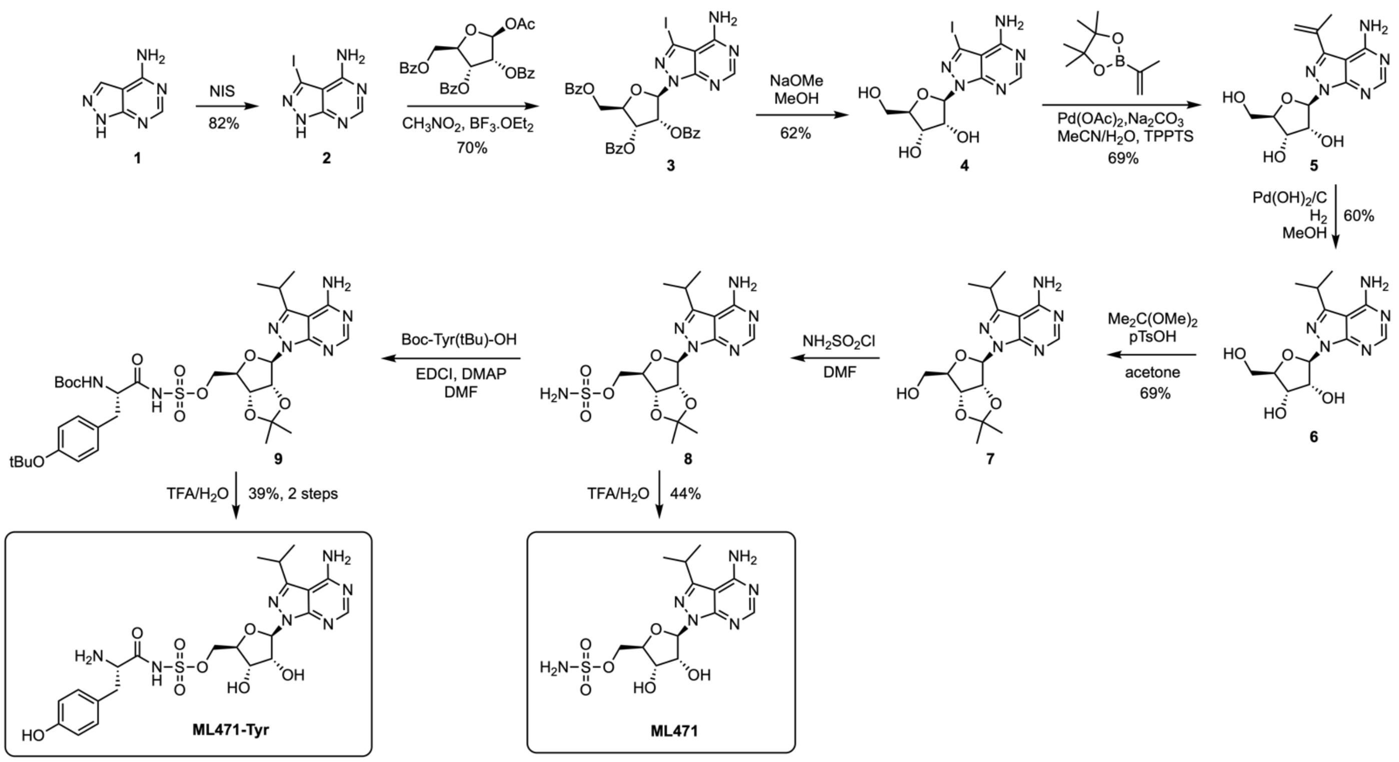

**Supplementary Figure 6. Synthetic routes to ML471 and Tyr-ML471.**

*3-iodo-1H-pyrazolo[3,4-d]pyrimidin-4-amine (**2**).*

4-Amino-1*H*-pyrazolo[3,4-*d*]pyrimidine (**1**, 5.0 g, 37 mmol) was dissolved in DMF (40 mL). *N*-Iodosuccinimide (NIS) (9.16 g, 40.7 mmol) was added, and the mixture was heated at 80 °C overnight. The mixture was cooled to room temperature and poured into ice-cold water (350 mL). The resulting suspension was stirred for 10 min at 0 °C then filtered. The solids were collected and dried *in vacuo* to afford **2** (8.8 g, 82% yield) as a white solid. $^1$H NMR (300 MHz, DMSO-$d_6$) δ 8.16 (1H, s), 13.80 (1H, br s) ppm. LC-MS: *m/z* 261.95 (M+H$^+$). Characterization data are in accordance with reported values [4].

*(2R,3R,4R,5R)-2-(4-amino-3-iodo-1H-pyrazolo[3,4-d]pyrimidin-1-yl)-5-((benzoyloxy)methyl)tetrahydrofuran-3,4-diyl dibenzoate (**3**).*

The pyrazolopyrimidine **2** (1.0 g, 3.83 mmol) and 1-*O*-acetyl-tri-*O*-benzoyl-β-D-ribofuranose (2.32 g, 4.6 mmol) were added to dry nitromethane (10 mL). The mixture was heated at reflux, then $BF_3 \cdot OEt_2$ (0.6 mL) was added, upon which the solids started to dissolve. After heating at reflux for 90 min, the solvent was removed *in vacuo*. The resulting oil was purified by silica column chromatography: excess 1-*O*-acetyl-tri-*O*-benzoyl-β-D-ribofuranose was eluted with 100% $CH_2Cl_2$, then elution with 5% acetone in $CH_2Cl_2$ afforded the product **3** (1.9 g, 70% yield) as a pale brown oil. $^1$H NMR (300 MHz, $CDCl_3$) δ 4.56–4.90 (3H, m), 6.20 (1H, t, *J* = 5.6 Hz), 6.36 (1H, dd, *J* = 5.0, 3.2 Hz), 6.76 (1H, d, *J* = 2.9 Hz), 7.35–7.62 (9H, m), 7.91–8.14 (6H, m), 8.34 (1H, s) ppm. LC-MS: *m/z* 706.07 (M+H$^+$). Characterization data are in accordance with reported values [4].

*(2R,3R,4S,5R)-2-(4-amino-3-iodo-1H-pyrazolo[3,4-d]pyrimidin-1-yl)-5-(hydroxymethyl)tetrahydrofuran-3,4-diol (**4**).*

Protected nucleoside **3** (1.15 g, 1.63 mmol) was dissolved in $CH_2Cl_2$ (0.85 mL). Methanol (8.5 mL) was added, followed by a solution of NaOMe in MeOH (4.5 M, 0.39 mL). The mixture was stirred at 60 °C for 2 h. The reaction was neutralised to pH 7 by the addition of 4 N HCl, and the solvents were removed *in vacuo*. The solid residue was purified by silica column chromatography (5–15% MeOH/$CH_2Cl_2$ gradient) to afford the final product **4** (0.400 g, 62% yield). $^1$H NMR (300 MHz, DMSO-$d_6$) δ 3.43 (1H, dd, *J* = 11.7, 5.6 Hz), 3.55 (1H, dd, *J* = 11.7, 4.4 Hz), 3.89 (1H, dd, *J* = 10.0, 4.4 Hz), 4.16 (1H, t, *J* = 4.7 Hz), 4.57 (1H, m), 6.03 (1H, d, *J* = 5.0 Hz), 8.23 (1H, s) ppm. LC-MS: *m/z* 394.00 (M+H$^+$). Characterization data are in accordance with reported values [4].

*(2R,3R,4S,5R)-2-(4-amino-3-(prop-1-en-2-yl)-1H-pyrazolo[3,4-d]pyrimidin-1-yl)-5-(hydroxymethyl)tetrahydrofuran-3,4-diol (**5**).*
Compound **4** (500 mg, 1.27 mmol) was added to 2-isopropenylboronic acid pinacol ester (317 mg, 3.81 mmol), $Na_2CO_3$ (405 mg 3.81 mmol), $Pd(OAc)_2$ (14.3 mg, 0.065 mmol), and TPPTS (87.4 mg, 0.15 mmol). Acetonitrile (2.5 mL) and $H_2O$ (5 mL) were added to the solids under nitrogen. After 5 min of stirring, the mixture was heated at reflux. Upon completion, the mixture was cooled to room temperature and neutralised to pH ~7 with 4 M aq. HCl. The mixture was concentrated *in vacuo* and the residue was purified by silica column chromatography (2–15% MeOH/$CH_2Cl_2$ gradient) to afford **5** (270 mg, 69% yield) as an off-white solid. $^1$H NMR (300 MHz, DMSO-$d_6$) δ 2.18 (3H, s), 3.45 (1H, m), 3.60 (1H, m), 3.92 (1H, dd, *J* = 9.7, 4.4 Hz), 4.23 (1H, dd, *J* = 9.7, 4.7 Hz), 4.59 (1H, dd, *J* = 9.4, 5.0 Hz), 4.84 (1H, t, *J* = 5.6 Hz), 5.12 (1H, d, *J* = 5.3 Hz), 5.29–5.45 (2H, m), 5.53 (1H, s), 6.13 (1H, d, *J* = 4.1 Hz), 8.23 (1H, s) ppm. LC-MS: *m/z* 308.14 (M+H$^+$). Characterization data are in accordance with reported values [4].

*(2R,3R,4S,5R)-2-(4-amino-3-isopropyl-1H-pyrazolo[3,4-d]pyrimidin-1-yl)-5-(hydroxymethyl)tetrahydrofuran-3,4-diol (**6**).*
Compound **5** (500 mg, 1.62 mmol) was dissolved in MeOH (10 mL) and $Pd(OH)_2$/C (50 mg) was added. The vessel was filled with $H_2$ at 20 psi, and the mixture was stirred overnight. The mixture was filtered through celite, and the solvents were removed *in vacuo*. The solid residue was purified by silica column chromatography (4–20% MeOH/$CH_2Cl_2$ gradient) to afford **6** (300 mg, 60% yield) as a white solid. $^1$H NMR (300 MHz, DMSO-$d_6$) δ 1.26 (6H, d, *J* = 6.7 Hz), 3.38–3.65 (3H, m), 3.90 (1H, dd, *J* = 9.4, 4.7 Hz), 4.25 (1H, dd, *J* = 10.3, 5.0 Hz), 4.56 (1H, dd, *J* = 10.0, 5.3 Hz), 4.85 (1H, dd, *J* = 6.7, 5.0 Hz), 5.07 (1H, d, *J* = 5.6 Hz), 5.32 (1H, d, *J* = 5.9 Hz), 6.06 (1H, d, *J* = 4.4 Hz), 8.15 (1H, s) ppm. LC-MS: *m/z* 310.14 (M+H$^+$). Characterization data are in accordance with reported values [4].

*((3aR,4R,6R,6aR)-6-(4-amino-3-isopropyl-1H-pyrazolo[3,4-d]pyrimidin-1-yl)-2,2-dimethyltetrahydrofuro[3,4-d][1,3]dioxol-4-yl)methanol (**7**).*
To compound **6** (570 mg, 1.84 mmol) in acetone (25 mL) was added 2,2-dimethoxypropane (1.13 mL, 9.2 mmol) and *p*-TsOH (350 mg, 1.84 mmol) and the mixture was stirred overnight at room temperature. Chilled aq. sat. $NaHCO_3$ was added and the mixture was stirred for 5 minutes, then dried *in vacuo*. The residue was dissolved by stirring in acetone for 1 h, the mixture was filtered and the filtrate evaporated *in vacuo*. The residue was purified by silica column chromatography (5% EtOH in EtOAc) to afford **7** (450 mg, 69% yield). $^1$H NMR (300 MHz, DMSO-$d_6$) δ 1.26 (6H, d, *J* = 6.7 Hz), 1.32 (3H, s), 1.51 (3H, s), 3.39 (1H, m), 3.50–3.62 (2H, m), 4.13 (1H, m), 4.94 (1H, d, *J* = 6 Hz), 5.23 (1H, m), 6.26 (1H, d, *J* = 1.8 Hz), 8.16 (1H, s) ppm. LC-MS: *m/z* 350 (M+H$^+$). Characterization data are in accordance with reported values [5].

*((3aR,4R,6R,6aR)-6-(4-amino-3-isopropyl-1H-pyrazolo[3,4-d]pyrimidin-1-yl)-2,2-dimethyltetrahydrofuro[3,4-d][1,3]dioxol-4-yl)methyl sulfamate (**8**).*
To compound **7** (425 mg, 1.21 mmol) in dry DMF (4.5 mL) was added sulfamoyl chloride (421 mg, 3.64 mmol) and the mixture was stirred at room temperature for 2 h. Several drops of methanol were added and the solvent was evaporated *in vacuo*. The compound was used in the next step without further purification. LC-MS: *m/z* 429.1 (M+H$^+$).

*((2R,3S,4R,5R)-5-(4-amino-3-isopropyl-1H-pyrazolo[3,4-d]pyrimidin-1-yl)-3,4-dihydroxytetrahydrofuran-2-yl)methyl sulfamate (**ML471**)*
Compound **8** (100 mg, 0.23 mmol) was dissolved in TFA/$H_2O$ (4:2). The mixture stirred at room temperature overnight, and completion of the deprotection was confirmed by LC-MS. The solvents were evaporated *in vacuo* and the residue was purified by HPLC (C8 column, 2–35% acetonitrile/$H_2O$) to afford the title

compound **ML471** (40 mg, 44% yield) as a white solid. $^1$H NMR (400 MHz, $CD_3OD$) δ 1.40 (6H, t, *J* = 7.0 Hz), 3.48 (1H, h, *J* = 6.8 Hz), 4.20–4.31 (2H, m), 4.35 (1H, dd, *J* = 10.2, 3.1 Hz), 4.63–4.68 (2H, m), 6.34 (1H, d, *J* = 1.9 Hz), 8.35 (1H, s). LC-MS: *m/z* 389.2 ($M+H^+$). Characterization data are in accordance with reported values [5].

*((3aR,4R,6R,6aR)-6-(4-amino-3-isopropyl-1H-pyrazolo[3,4-d]pyrimidin-1-yl)-2,2-dimethyltetrahydrofuro[3,4-d][1,3]dioxol-4-yl)methyl ((S)-3-(4-(tert-butoxy)phenyl)-2-((tert-butoxycarbonyl)amino)propanoyl)sulfamate (**9**).*

Compound **8** (50 mg, 0.12 mmol) was added to Boc-Tyr(*t*Bu)-OH (60.7 mg, 0.18 mmol), EDCI (92.0 mg, 0.48 mmol), DMAP (16.1 mg, 0.13 mmol) in dry DMF (5 mL) and the mixture was stirred overnight at room temperature. The solvent was evaporated *in vacuo*. The residue was added to water and the organic layer was extracted using EtOAc, dried using $Na_2SO_4$, and evaporated *in vacuo* to give a crude sample of **9** (100 mg), a viscous pale brown oil, which was used in the next step without further purification. LC-MS: *m/z* 748.1 ($M+H^+$).

*((2R,3S,4R,5R)-5-(4-amino-3-isopropyl-1H-pyrazolo[3,4-d]pyrimidin-1-yl)-3,4-dihydroxytetrahydrofuran-2-yl)methyl (L-tyrosyl)sulfamate (**Tyr-ML471**)*

The crude compound **9** (100 mg) was dissolved in TFA (0.8 mL) and $CH_2Cl_2$ (4 mL) and the mixture was stirred at room temperature for 2 h. LC-MS confirmed deprotection of the tBu and Boc groups (*m/z* 592 (M+H)). Water (0.4 mL) was added, and the mixture was stirred at room temperature overnight. The volatiles were removed *in vacuo* then the residue was purified by column chromatography (5–15% MeOH/$CH_2Cl_2$ gradient), followed by HPLC purification (C8 column, 2–35% acetonitrile/$H_2O$) to afford the title compound **Tyr-ML471** (25 mg, 34% yield over 2 steps) as an off-white solid. $^1$H NMR (400 MHz, DMSO-*d*$_6$) 1.26 (6H, d, *J* = 6.7 Hz), 2.64–2.88 (2H, m), 3.02 (1H, m), 3.98–4.12 (2H, m), 4.24 (1H, m), 4.37 (1H, t, *J* = 5.1 Hz), 4.51 (1H, m), 6.13 (1H, d, *J* = 3.0 Hz), 6.64–6.74 (2H, m), 7.01–7.05 (2H, m), 8.27 (1H, s). LC-MS: *m/z* 552.2 ($M+H^+$).

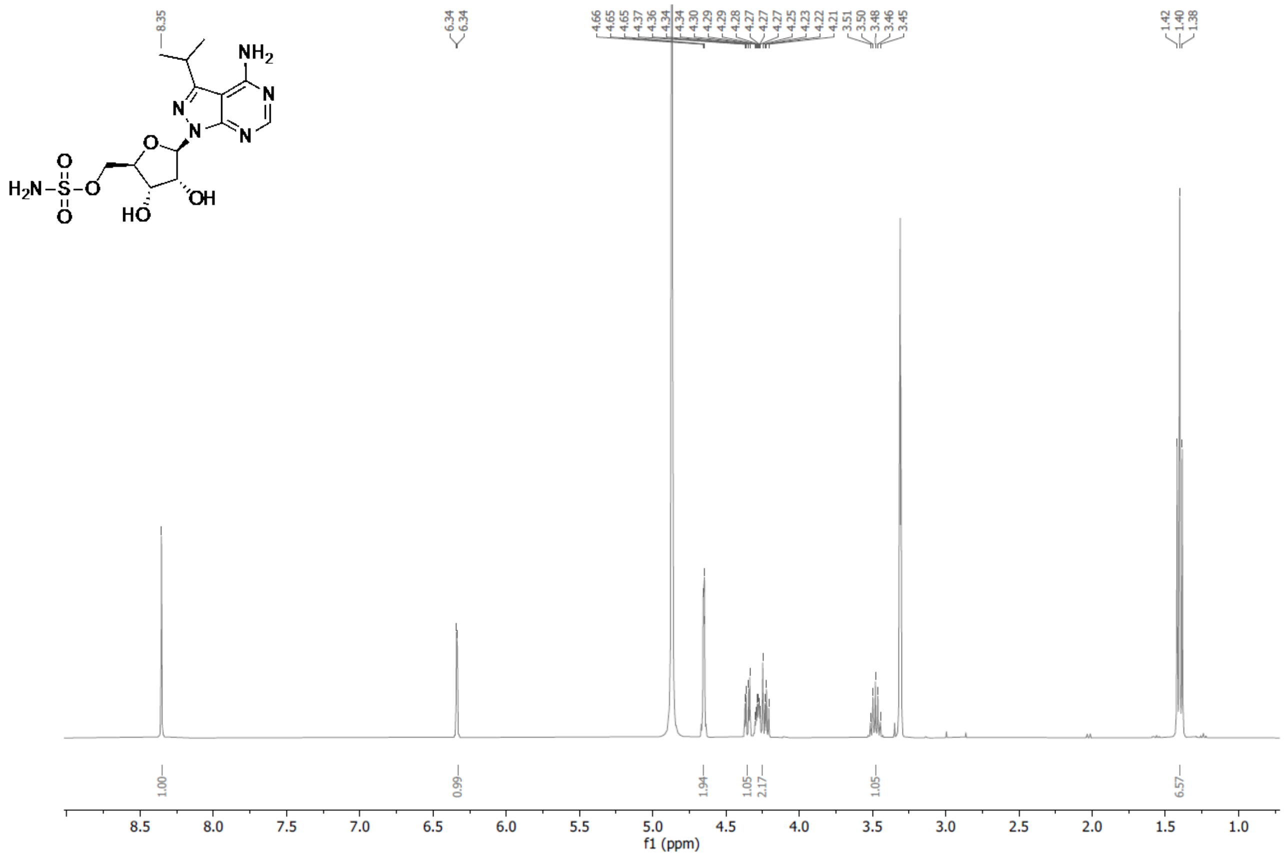

**Supplementary Figure 7. $^1$H NMR Spectrum of ML471**

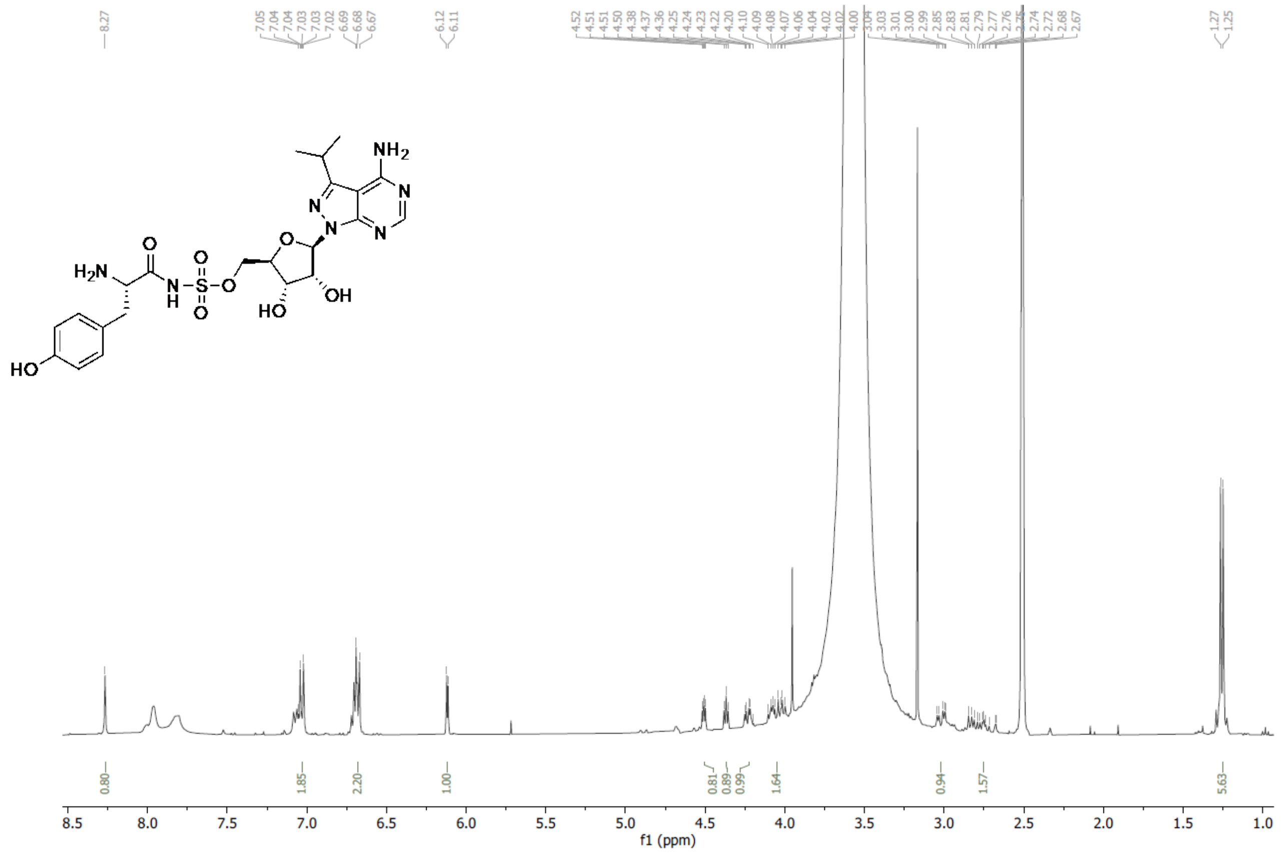

**Supplementary Figure 8. $^1$H NMR Spectrum of Tyr-ML471**

### References

1. Xie SC, Metcalfe RD, Dunn E, Morton CJ, Huang SC, Puhalovich T, et al. Reaction hijacking of tyrosine tRNA synthetase as a new whole-of-life-cycle antimalarial strategy. Science. 2022;376(6597):1074-9.
2. Baragaña B, Hallyburton I, Lee MCS, Norcross NR, Grimaldi R, Otto TD, et al. A novel multiple-stage antimalarial agent that inhibits protein synthesis. Nature. 2015;522(7556):315-20.
3. Sanz LM, Crespo B, De-Cozar C, Ding XC, Llergo JL, Burrows JN, et al. *P. falciparum* in vitro killing rates allow to discriminate between different antimalarial mode-of-action. PloS one. 2012;7(2):e30949.
4. Bouton J, Ferreira de Almeida Fiuza L, Cardoso Santos C, Mazzarella MA, Soeiro MdNC, Maes L, et al. Revisiting pyrazolo[3,4-d]pyrimidine nucleosides as anti-*Trypanosoma cruz*i and antileishmanial agents. Journal of medicinal chemistry. 2021;64(7):4206-38.
5. Adhikari S, Calderwood EF, England DB, Gould AE, Harrison SJ, Huang S-C, et al. Atg7 inhibitors and the uses thereof. World Intellectual Property Organization. 2017;WO/2018/089786 PCT/US2017/061094.
